# Global distributions and plant associations of the ecologically diverse ascomycete order Helotiales

**DOI:** 10.64898/2026.09.28.754902

**Authors:** Hirokazu Toju, Aryan Sathish Raj, Mikihito Noguchi, Akifumi Murata

## Abstract

**Background:** Fungi in plant root systems are key to understanding the dynamics and functioning of terrestrial ecosystems. Whereas our knowledge of below-ground plant–fungus symbioses has derived mainly from studies of mycorrhizal and pathogenic fungi, recent studies have uncovered an unexpected diversity of ecological niches occupied by root-associated fungi. The fungal order Helotiales is conspicuous in this respect because this predominantly root-associated taxon encompasses a broad ecological spectrum, ranging from saprotrophic and pathogenic fungi to endophytic and mycorrhizal fungi with plant growth-promoting functions. In this study, we integrated a worldwide dataset of 1,996 Helotiales Species Hypotheses (i.e., species-level taxa tentatively defined based on DNA sequences) detected in roots of 362 plant genera representing 117 families and 47 orders, and systematically quantified the specificity of Helotiales–plant associations.

**Results:** The analysis of 37,688 unique occurrence records of Helotiales fungi across 1,940 global sampling locations revealed highly structured associations between Helotiales genera and plant lineages, exemplified by the strong specificity of *Hyaloscypha*, *Oidiodendron*, and *Phialocephala* for Pinales and that of *Glutinomyces* for Fagales, even after statistically controlling background biogeographic patterns. At the Species-Hypothesis-level analysis, major clades within the Helotiales phylogeny consistently exhibited high specificity for Pinaceae, whereas highly specific associations with other ectomycorrhizal, arbuscular mycorrhizal, and nonmycorrhizal plants were restricted to small terminal clades of the fungal phylogeny.

**Conclusions:** The integration of ecological specificity and phylogenetic analyses suggests the possibility that Helotiales fungi established close associations with Pinaceae early in their evolutionary history and subsequently underwent host shifts to diverse angiosperm lineages. Given the emerging evidence that Helotiales fungi play pivotal ecological and physiological roles in plant root systems, these statistical insights into Helotiales–plant associations provide a fundamental basis for understanding the evolutionary history and functioning of terrestrial ecosystems.

## BACKGROUND

The order Helotiales is one of the most species-rich and ecologically diverse lineages within the fungal phylum Ascomycota [1–5]. Its members occupy a broad range of ecological niches as saprotrophs, plant pathogens, aquatic decomposers, root and foliar endophytes, and ectomycorrhizal and ericoid mycorrhizal symbionts [6–9]. Despite their potentially important roles in terrestrial and aquatic ecosystem processes, the diversity and ecological functions of Helotiales have long been underestimated [10,11], partly because many species produce tiny fruiting bodies, remain sterile in culture, or are known only from environmental DNA sequences [12,13]. Nonetheless, technical advances in DNA sequencing have begun to reshape our understanding of the diversity and ecology of Helotiales fungi [14,15]. Amplicon sequencing approaches, for example, have revealed extensive cryptic diversity of Helotiales especially in plant root systems [16–19]. Moreover, genomic and transcriptomic approaches have begun to reveal unique combinations of saprotrophic and symbiotic lifestyles within the fungal order [20–22], which makes a clear contrast with the convergent losses of saprotrophic functions in most mycorrhizal clades [23].

Recent inoculation experiments and molecular studies have further shown that ecological functions can vary considerably even among closely related Helotiales fungi [24,25]. For example, lineages related to ericoid mycorrhizal fungi (e.g., *Hyaloscypha hepaticicola*) may form ectomycorrhizas or ectomycorrhiza-like associations with non-ericaceous hosts, indicating substantial ecological and evolutionary flexibility in their root-association strategies [25–27]. In addition, Helotiales include “dark septate endophytes” that colonize roots with melanized septate hyphae and microsclerotia [4,9,28,29]. Such endophytic fungi form structures within roots that are morphologically distinct from conventional mycorrhizas, and they can associate with diverse plant taxa traditionally recognized as nonmycorrhizal (e.g., Brassicaceae) [13,14]. Many of these endophytic fungi play important physiological and ecological roles in plant root systems by provisioning nitrogen and other nutrients [11,13,28,30–32]. They can also enhance plant resistance to acidic and metal-contaminated soils [24,33] as well as resistance against pests and pathogens [34–36]. Moreover, Helotiales endophytic fungi often coexist with mycorrhizal fungi in plant root tips [37], potentially influencing the assembly of entire root-associated mycobiomes in host-specific ways [38]. Thus, although still underappreciated, the diversity and ecology of Helotiales fungi are key to understanding the dynamics and functioning of terrestrial ecosystem processes.

Given the potential prevalence and ecosystem functions of plant-associated Helotiales fungi, it is crucial to obtain an overview of their diversity across a broad range of associated plant lineages. The diversity of root-associated Helotiales has been explored mainly in studies targeting ericoid mycorrhizal (Ericaceae) and ectomycorrhizal (e.g., Pinaceae, Fagaceae, and Betulaceae) woody plants [3,7,16,39]. Nonetheless, Helotiales fungi have also been detected in the roots of diverse nonmycorrhizal and arbuscular mycorrhizal plants in forest and grassland ecosystems spanning arctic to tropical regions [10,15,17,40]. In particular, Helotiales endophytes can promote the growth of Brassicaceae and Poaceae plants by enhancing nitrogen or phosphorus acquisition from nutrient-poor soils in alpine habitats and heathlands [11,13,41]. Furthermore, inoculation experiments have shown that Helotiales isolates can promote plant growth and enhance resistance to pathogens or herbivores in diverse herbaceous plants, including tropical-origin crops such as rice and banana [34,36]. These findings indicate that the diversity and physiological properties of Helotiales–plant associations remain incompletely understood. Although Helotiales endophytes may have been overlooked in studies focusing primarily on mycorrhizal symbioses, their presence in plant roots can be detected retrospectively from DNA sequence data [7,12]. Thus, compiling DNA sequence records and associated metadata from public databases provides an opportunity to systematically evaluate the global prevalence and compositions of Helotiales–plant associations.

In this study, we analyzed a worldwide dataset of fungal DNA sequences to systematically evaluate the specificity of associations between Helotiales fungi and diverse plant lineages. We screened the GlobalFungi database [42] for Helotiales records with associated plant metadata, thereby constructing a large data matrix comprising 1,996 Helotiales Species Hypotheses (SHs) and 362 plant genera. First, we evaluated among-continent variation in the occurrence records of individual Helotiales lineages to assess the potential contributions of background biogeographic patterns to associations with plants. We then quantified the specificity of Helotiales–plant associations using a randomization approach that statistically controlled for these background biogeographic patterns. By retrieving nucleotide sequence information for individual SHs from the UNITE database [43], we further evaluated the phylogenetic conservatism or lability of plant-association specificity across Helotiales lineages. Overall, this study provides a fundamental ecological perspective on a ubiquitous plant-associated fungal lineage and establishes a platform for understanding community- and ecosystem-scale phenomena that cannot be explained solely from a mycorrhiza-centric view of below-ground plant–fungus symbioses.

## MATERIALS AND METHODS

### Database

We reconstructed the global distribution of plant-associated fungi in the order Helotiales using the GlobalFungi database [42]. We downloaded release 5.0 of the GlobalFungi dataset (https://globalfungi.com/), including fungal internal transcribed spacer 1 and 2 (ITS1 and ITS2) sequence records and their associated sample metadata (GlobalFungi_5_SH_abundance_ITS1_ITS2.txt.gz and GlobalFungi_5_sample_metadata.txt.gz, respectively). By screening the occurrence data, we identified 3,396 unique Species Hypotheses (SHs), defined as DNA sequence-based species-level groupings used in the UNITE database [43], that were assigned to the order Helotiales. In total, the dataset contained 1,309,226 Helotiales occurrence records from 19,372 sampling locations (sampling sites) across 127 countries, with sampling locations defined as unique combinations of latitude and longitude (Fig. S1).

To analyze the plant associations of Helotiales fungi, we focused on occurrence records for which the “sample_type” field in the metadata was designated as “root”. We then further screened these records to retain only those for which a single plant name was specified in either the “host_or_vegetation” or “dominant_plant_species” column of the metadata file. When information was available in both columns, the “host_or_vegetation” entry was prioritized over the “dominant_plant_species” entry. This screening yielded 37,886 records representing unique combinations of Helotiales SH, associated plant identity, and sampling location (hereafter referred to as “unique occurrences”) from 1,941 sampling locations across 41 countries. After removing 198 unique occurrences with ambiguous plant taxonomic assignments, the final dataset comprised 37,688 unique occurrences from 1,940 sampling locations, including 1,996 Helotiales SHs representing 184 genera and 36 families and 362 plant genera representing 117 families and 47 orders (Fig. 1; Figs. S1–4). For fungal taxonomy, synonymous names, such as *Hyaloscypha* and *Meliniomyces*, were retained as recorded in the GlobalFungi database.

**Figure 1:**
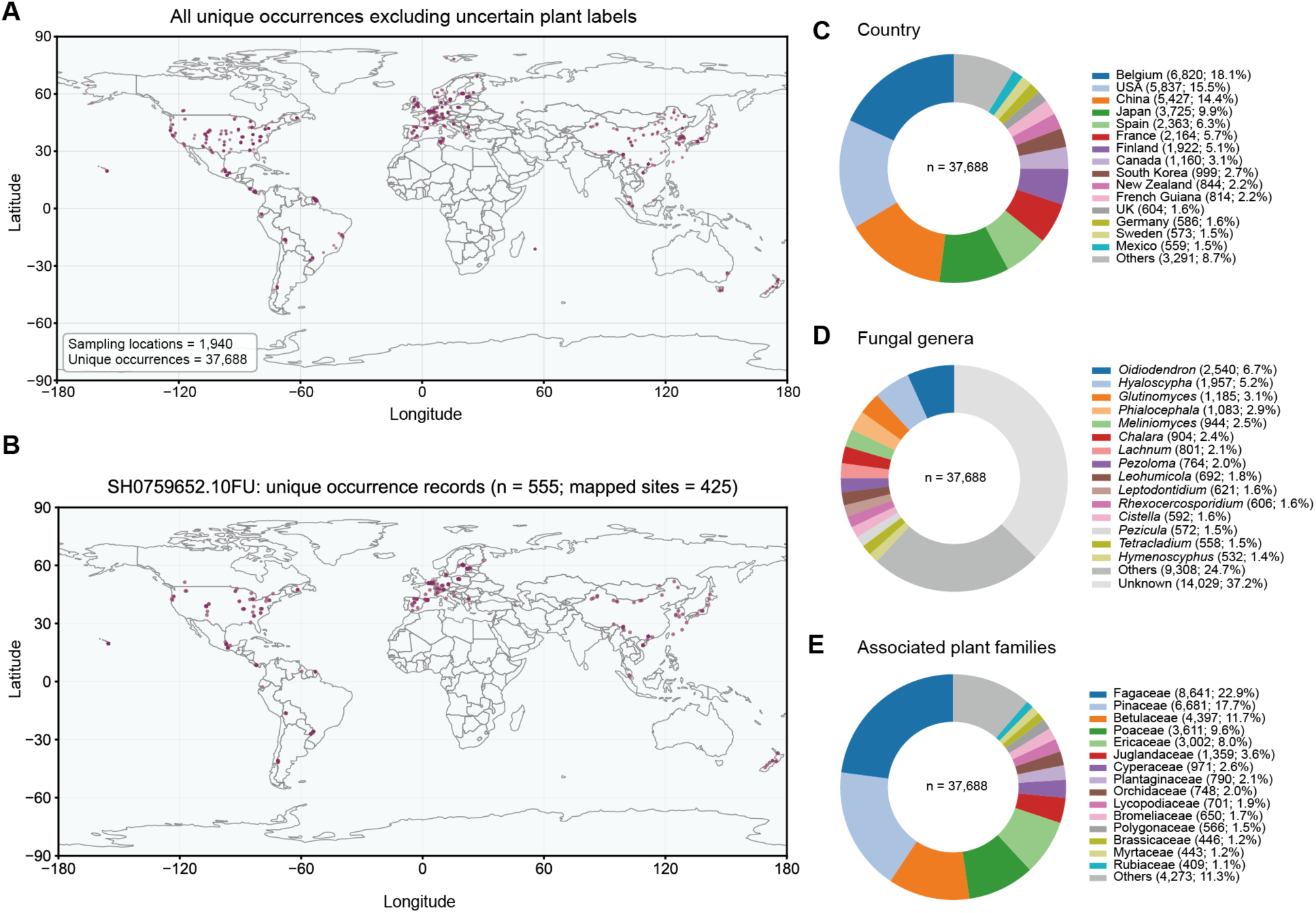
Overview of the dataset. (A) Map of sampling locations for which Helotiales fungi were recorded with associated plant information. (B) Map of sampling locations at which the most frequently observed Helotiales SH (SH0759652.10FU) was detected with associated plant information. (C) Composition of unique Helotiales occurrence records by source country. (D) Composition of unique Helotiales occurrence records by fungal genus. (E) Composition of unique Helotiales occurrence records by associated plant family.

Within the screened data, unique occurrences from Europe (42.9%), Asia (27.2%), and North America (20.8%) predominated, although the dataset also included records from South America (3.7%), Australia (2.5%), and Africa (2.5%) (Fig. 1C; Fig. S2). The most frequently observed fungal SH was represented by 555 unique occurrences, whereas the 50th most frequently observed SH was represented by 137 unique occurrences (Fig. 1B; Fig. S3). SHs assigned to *Oidiodendron* (6.7%), *Hyaloscypha* (5.2%), *Glutinomyces* (3.1%), *Phialocephala* (2.9%), and *Meliniomyces* (2.5%) collectively accounted for 20.5% of the dataset (Fig. 1D). The associated plants included families predominantly forming ectomycorrhizal associations, such as Fagaceae, Pinaceae, Betulaceae, Salicaceae, and Myrtaceae (Fig. 1E; Fig. S4). The dataset also included the ericoid mycorrhizal family Ericaceae and predominantly arbuscular mycorrhizal families such as Poaceae, Plantaginaceae, Asteraceae, and Rosaceae. In addition, it included the orchid mycorrhizal family Orchidaceae and predominantly nonmycorrhizal families such as Brassicaceae, Cyperaceae, and Polygonaceae.

### Geographic distributions

To evaluate how Helotiales lineages differ in their geographic distributions, we performed a randomization analysis. The specificity of each Helotiales fungal genus for each geographic region (defined by the “continent” column of the GlobalFungi metadata) was quantified based on the *d*′ metric of specificity [44]. We then standardized the *d*′ specificity index based on a randomization approach in which the geographic region label was shuffled across occurrences in the data matrix (10,000 permutations). Specifically, for each fungal genus, a *z*-standardized *d*′ score was obtained as follows:

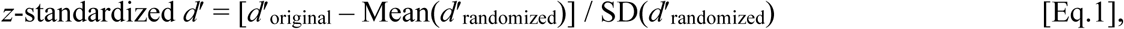

where *d*′_original_ was the *d’* estimate of the original data, and Mean(*d*′_randomized_) and SD(*d*′_randomized_) were the mean and standard deviation of the *d’* scores of randomized data matrices. The relationship between the *z*-standardized *d*′ scores and *P* values was examined after the Benjamini–Hochberg correction [false discovery rates (FDR)]. The “bipartite” 2.24 package [45] of R 4.6.0 [46] was used in the calculation of *d*′ scores. Moreover, specificity of each fungus–geographic region association was evaluated using the “two-dimensional preference (2DP)” metric [16]. Specifically, the extent to which focal fungus–geographic region associations were observed more frequently than that expected by chance was calculated for a fungal genus *i* and geographic region *j* as follows:

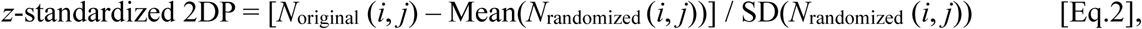

where *N*_original_ (*i*, *j*) denoted the number of unique occurrences from which a focal combination of a fungal genus and a geographic region was observed in the original data, and the Mean(*N*_randomized_ (*i*, *j*)) and SD(*N*_randomized_(*i*, *j*)) were the mean and standard deviation of the number of unique occurrences for the focal fungus–geographic region pair across randomized matrices (10,000 permutations). The top-50 most frequently observed fungal genera were targeted in the analysis.

### Specificity in Helotiales–plant associations

We next quantified the specificity of associations between Helotiales fungi and plant lineages. Using the *z*-standardized *d′* specificity metric (Eq. 1), we calculated the specificity of fungal associations with plant orders for the 50 most frequently observed Helotiales genera. Occurrence records involving the 30 most frequently observed plant orders were included in this analysis. To account for the geographic distributions of plant taxa, plant labels were shuffled among occurrence records within spatial blocks defined by 20° latitudinal intervals from the Northern to the Southern Hemisphere in the randomization analysis (10,000 permutations). An analysis using an alternative definition of spatial blocks based on the “continent” column of the GlobalFungi metadata (Fig. S2) yielded qualitatively and quantitatively similar results. We also quantified the specificity of plant orders for their associations with Helotiales genera using the *z*-standardized *d′* metric. In addition, we calculated a *z*-standardized 2DP score (Eq. 2) for each combination of fungal genus and plant order using the same spatially blocked randomization procedure (10,000 permutations). To evaluate association specificity at a finer taxonomic resolution, we conducted an additional analysis focusing on the 80 most frequently observed fungal SHs and the 30 most frequently observed plant families.

### Phylogenetic signals in Helotiales–plant associations

Based on the specificity analyses described above, we examined the phylogenetic conservatism or evolutionary lability of Helotiales fungi in their associations with plant lineages. We first expanded the analysis of Helotiales–plant associations to include the 150 most frequently observed Helotiales SHs. Among the 150 SHs, nucleotide sequences covering the ITS1, 5.8S rRNA, and ITS2 regions were available for 98 SHs in the UNITE 04.04.2024 release database (https://unite.ut.ee/repository.php). The sequences were aligned using the L-INS-i algorithm implemented in MAFFT v7.490 [47], with the Leotiales fungus *Microglossum griseoviride* (SH0748249) designated as the outgroup. Unambiguously aligned nucleotide sites were then selected using trimAl v1.5.rev1 [48]. A neighbor-joining tree was constructed under the Kimura two-parameter substitution model [49] using the R package ape v5.8.1 [50], with branch support evaluated using 1,000 bootstrap replicates. Phylogenetic conservatism in the *z*-standardized *d′* scores of plant-association specificities across fungal SHs was evaluated using Pagel’s *λ* [51], as implemented in the R package phytools v2.5.2 [52]. To assess the robustness of the results, the analysis of Pagel’s *λ* was repeated using a maximum-likelihood phylogeny reconstructed with IQ-TREE v3.1.2 [53], for which the TIM2+F+I+G4 substitution model [54] was selected.

In addition to the analysis examining overall specificity for plants, we tested the phylogenetic conservatism in associations with specific plant lineages. In this respect, the *z*-standardized 2DP scores for each plant family were examined based on the Pagel’s *λ* statistic.

### Use of generative AI tools

ChatGPT 5.5 (OpenAI) was used to assist in developing the R (v.4.6.0) and Python (v3.9.6) scripts used for data screening, randomization analyses, and figure generation. All AI-assisted code was reviewed, tested, and validated by the authors, who take full responsibility for the analyses and their results. Generative AI tools were not used to generate or interpret data, or to draw scientific conclusions.

## RESULTS

### Geographic distributions

Genera in the order Helotiales showed remarkable variation in their global distributions (Fig. 2). *Hyaloscypha* and its synonym *Meliniomyces* were both significantly associated with Europe (*z*-standardized 2DP = 9.63 and 16.28, respectively; FDR-adjusted *P* < 0.001 for both). Likewise, several other genera, including *Pezicula*, *Rhizodermea*, *Chalara*, *Hymenoscyphus*, *Mollisia*, *Tricladium*, *Coleophoma*, *Mycoarthris*, *Lachnellula*, *Sclerotinia*, *Endoradiciella*, and *Amicodisca*, showed significantly Europe-biased distributions (FDR-adjusted *P* < 0.001 for all). In contrast, among the five most frequently observed genera, *Glutinomyces* showed a strong association with Asia (*z*-standardized 2DP = 11.38; FDR-adjusted *P* < 0.001). Other genera showing significantly Asia-biased distributions included *Cryptosporiopsis*, *Hyphodiscus*, *Fontanospora*, and *Gyoerffyella* (FDR-adjusted *P* < 0.001 for all). The analysis further showed that *Lachnum*, *Cistella*, *Tetracladium*, *Spirosphaera*, *Clathrosphaerina*, *Microscypha*, and *Gremmenia* exhibited significantly North America-biased distributions (FDR-adjusted *P* < 0.001 for all). South America was particularly strongly associated with *Pseudohelotium* (*z*-standardized 2DP = 31.73; FDR-adjusted *P* < 0.001) and *Xylogone* (*z*-standardized 2DP = 14.07; FDR-adjusted *P* < 0.001).

**Figure 2:**
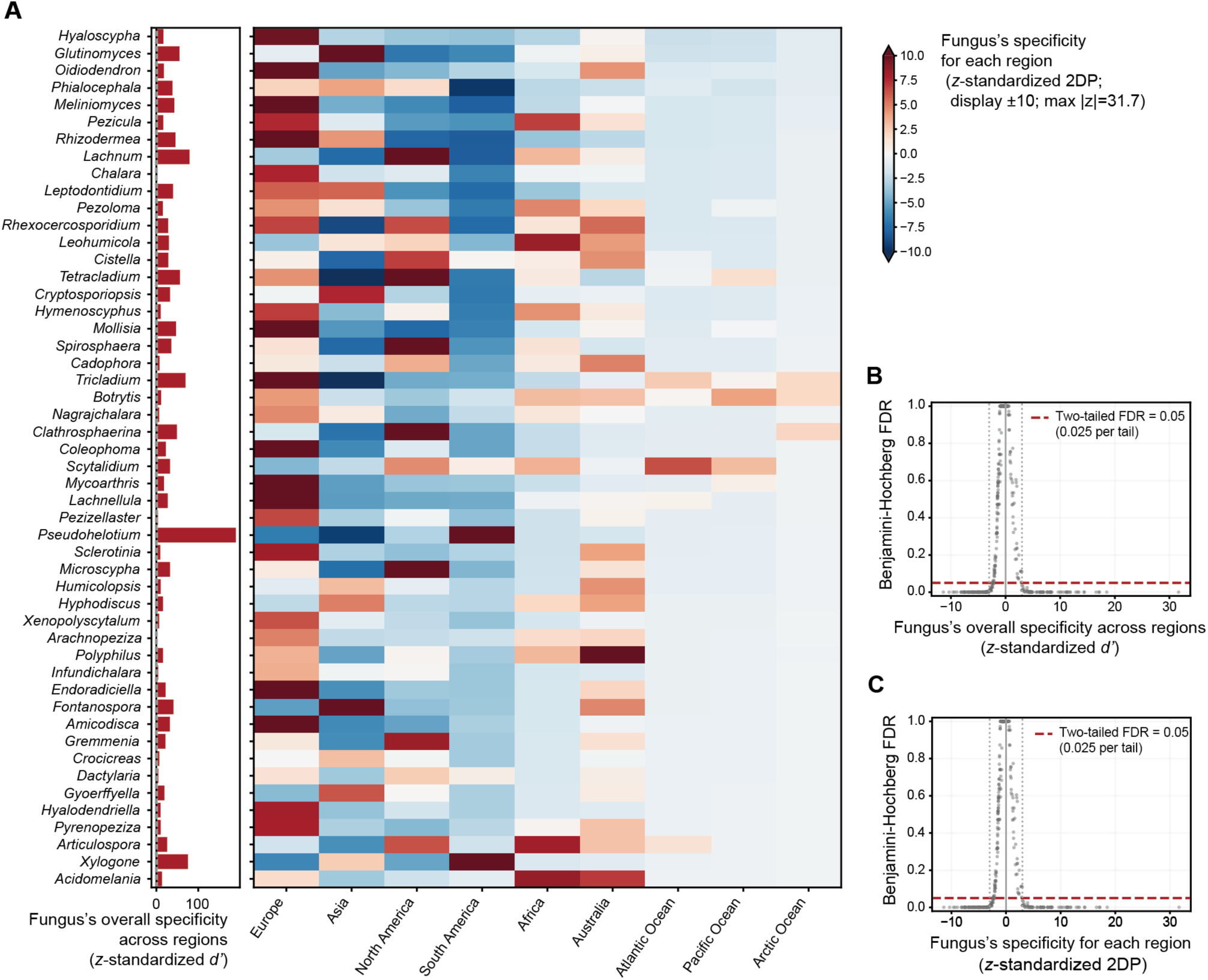
Biogeographic patterns in the dataset. (A) Specificity of each Helotiales genus for geographic regions. For each Helotiales genus, specificity across regions was quantified using the *z*-standardized *d′* metric (left). Specificity for individual regions was also evaluated using the *z*-standardized 2DP metric (right). Given that a *z*-standardized 2DP value larger than 3 or smaller than −3 roughly indicates significant specificity (panel C), heatmap colors are scaled between −10 and 10 to emphasize variation around the significance thresholds. (B) Statistical significance of the *z*-standardized *d′* metric. A *z*-standardized *d′* value larger than 3 or smaller than −3 roughly indicates significant specificity after Benjamini–Hochberg adjustment of *P*-values, with a two-tailed FDR threshold of 0.05 (0.025 per tail). (C) Statistical significance of the *z*-standardized 2DP metric.

Although fewer occurrence records were available from the remaining biogeographic regions than from Europe, Asia, North America, and South America, several Helotiales genera showed significantly biased distributions in these regions (Fig. 2). For example, *Leohumicola*, *Articulospora*, and *Acidomelania* were overrepresented in Africa (FDR-adjusted *P* < 0.001 for all), whereas *Polyphilus* showed significantly Australia-biased distributions (FDR-adjusted *P* < 0.01 for all). *Scytalidium* was also overrepresented in the Atlantic Ocean (FDR-adjusted *P* < 0.001).

### Specificity in Helotiales–plant associations

The randomization analysis, which controlled for biased fungal distributions across biogeographic regions, revealed highly specialized associations between Helotiales fungi and plant lineages (Fig. 3). Among the five most frequently observed Helotiales genera, *Glutinomyces* showed significantly high association specificity toward Fagales (*z*-standardized 2DP = 22.69; FDR-adjusted *P* < 0.01). Other fungal genera significantly associated with Fagales included *Rhizodermea*, *Tricladium*, *Coleophoma*, and *Amicodisca* (FDR-adjusted *P* < 0.01 for all). Many genera also exhibited their strongest plant-order associations with Pinales. *Hyaloscypha* and its synonym *Meliniomyces*, as well as *Oidiodendron*, *Phialocephala*, *Chalara*, *Leptodontidium*, *Pezoloma*, *Leohumicola*, *Cryptosporiopsis*, *Cadophora*, *Pseudohelotium*, *Humicolopsis*, *Hyphodiscus*, *Xenopolyscytalum*, *Infundichalara*, and *Crocicreas*, showed their highest *z*-standardized 2DP scores for Pinales and were significantly overrepresented in roots of Pinales plants (FDR-adjusted *P* < 0.01 for all). Meanwhile, *Mollisia*, *Fontanospora*, and *Pyrenopeziza* showed their highest *z*-standardized 2DP scores for Ericales and were significantly overrepresented in roots of Ericales plants (FDR-adjusted *P* < 0.01 for all).

**Figure 3:**
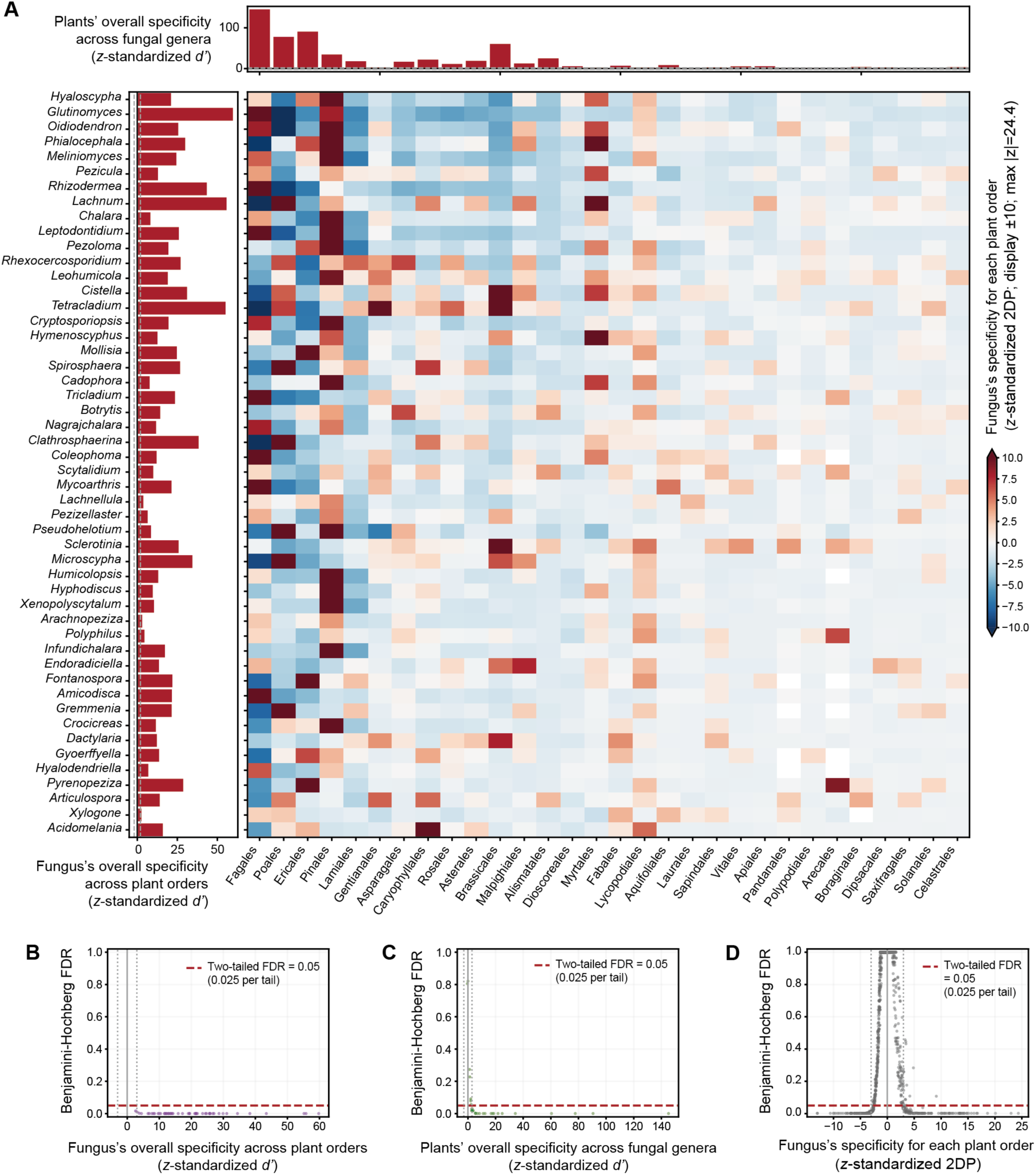
Specificity of associations between Helotiales genera and plant orders. (A) Specificity of each Helotiales genus for plant orders. For each Helotiales genus, specificity across plant orders was quantified using the *z*-standardized *d′* metric (left). Specificity for individual plant orders was also evaluated using the *z*-standardized 2DP metric (heatmap). Overall specificity of each plant order across Helotiales genera was quantified using the *z*-standardized *d′* metric (top). Given that a *z*-standardized 2DP value larger than 3 or smaller than −3 roughly indicates significant specificity (panel D), heatmap colors are scaled between −10 and 10 to emphasize variation around the significance thresholds. (B) Statistical significance of the *z*-standardized *d′* metric for Helotiales genera. (C) Statistical significance of the *z*-standardized *d′* metric for plant orders. (D) Statistical significance of the *z*-standardized 2DP metric.

Beyond the ectomycorrhizal and ericoid mycorrhizal plant lineages commonly examined as hosts of root-associated Helotiales fungi, diverse plant orders showed preferential associations with specific fungal genera (Fig. 3). For example, *Lachnum*, *Clathrosphaerina*, and *Microscypha* showed very high *z*-standardized 2DP scores for Poales and were significantly overrepresented in roots of Poales plants (FDR-adjusted *P* < 0.01 for all). Likewise, *Cistella*, *Tetracladium*, *Sclerotinia*, and *Dactylaria* showed their strongest plant-order associations with Brassicales (FDR-adjusted *P* < 0.01 for all). Highly specific associations were also observed for *Rhexocercosporidium* and *Botrytis* with Asparagales; *Acidomelania* with Caryophyllales; *Endoradiciella* with Malpighiales; *Phialocephala*, *Lachnum*, and *Hymenoscyphus* with Myrtales; *Pyrenopeziza* with Arecales.

Although the genus-level associations with plant orders were largely reflected in the SH-level analysis of associations with plant families, the finer taxonomic resolution provided additional insights into Helotiales–plant associations (Fig. 4; Fig. S5). For example, individual fungal SHs showed distinct association specificities with plant families within the order Fagales, including Fagaceae, Betulaceae, Juglandaceae, and Nothofagaceae. Likewise, the association specificity of individual SHs differed between Poaceae and Cyperaceae within the order Poales. Many fungal SHs showed strong associations with specific plant families. Examples included SH0851808, SH0907977, and SH0948790 associated with Bromeliaceae; SH0908345 and SH1012022 with Rubiaceae; SH0963754 with Brassicaceae.

**Figure 4:**
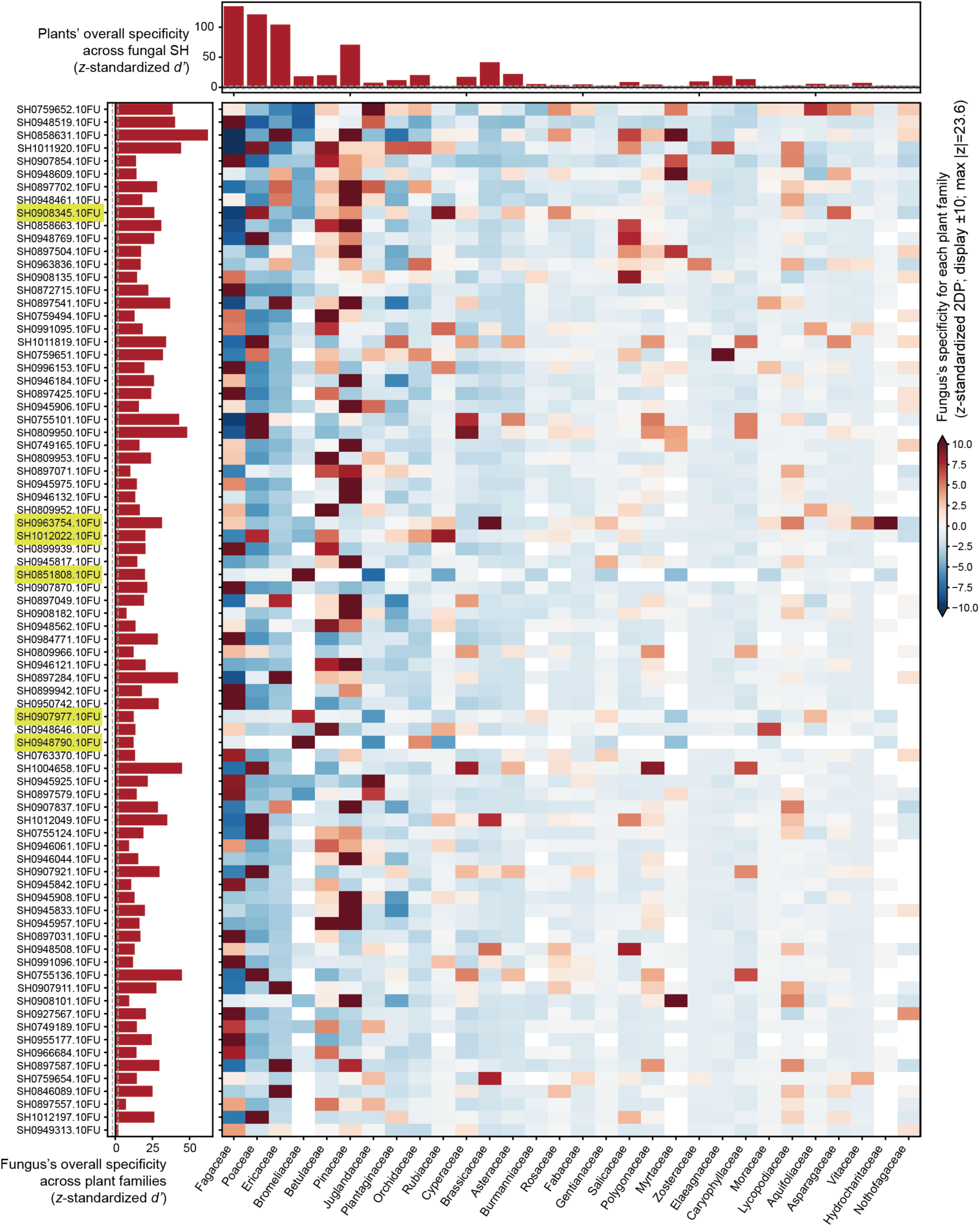
Specificity of associations between Helotiales SHs and plant families. Specificity of each Helotiales SH for plant families was quantified using the *z*-standardized *d′* metric (left). Specificity for individual plant families was also evaluated using the *z*-standardized 2DP metric (heatmap). Overall specificity of each plant family across Helotiales SHs was quantified using the *z*-standardized *d′* metric (top). Given that a *z*-standardized 2DP value larger than 3 or smaller than −3 roughly indicates significant specificity (Fig. S5), heatmap colors are scaled between −10 and 10 to emphasize variation around the significance thresholds. SH IDs mentioned in the main text are highlighted in yellow.

### Phylogenetic signals in Helotiales–plant associations

The *z*-standardized *d′* scores representing the overall specificity of Helotiales fungi for plant associations were consistently high throughout the Helotiales phylogeny (Figs. S6–7). As expected from this limited variation, no significant phylogenetic signal was detected in the overall degree of association specificity toward plant families in either the neighbor-joining analysis (Pagel’s *λ* = 0.318, *P* = 0.3397) or the maximum-likelihood analysis (Pagel’s *λ* = 0.235, *P* = 0.2847). Likewise, when associations with individual plant families were examined, specificity toward Fagaceae and Ericaceae, for example, showed no significant phylogenetic signal (Table S1). Nonetheless, significant phylogenetic signals were detected for associations with Poaceae, Pinaceae, Orchidaceae, Fabaceae, Salicaceae, Caryophyllaceae, Cactaceae, and Vitaceae in both the neighbor-joining and maximum-likelihood analyses (Table S1).

Conspicuous phylogenetic signals were detected for specificity toward Pinaceae. Closely related SHs within the *Oidiodendron, Phialocephala*, and *Hyaloscypha* (*Meliniomyces*) clades consistently exhibited high specificity toward this plant family, whereas the *Leptodontidium*– *Glutinomyces* clade comprised SHs with varying degrees of association specificity with Pinaceae (Fig. 5). Along with these clades including SHs with exceptionally strong associations with Pinaceae, most other clades within the Helotiales phylogeny showed positive association specificity toward this plant lineage (Fig. 5). In contrast, positive associations with other ectomycorrhizal plant families, such as Fagaceae (Fig. 6), Betulaceae (Fig. S8), Salicaceae (Fig. S9), and Myrtaceae (Fig. S10), were observed only in a few small terminal clades within the Helotiales phylogeny. Likewise, positive associations with the ericoid mycorrhizal plant family Ericaceae were confined to small terminal clades or individual SHs (Fig. 7). Although the strength of phylogenetic conservatism varied among target plant families (Table S1), similar phylogenetically structured patterns of association specificity were also observed in analyses targeting arbuscular mycorrhizal (Figs. S11– 26), orchid mycorrhizal (Fig. S27), and nonmycorrhizal plant families (Figs. S28–33). For example, conserved high specificity toward Poaceae was observed within the predominantly saprotrophic genus *Lachnum* and a poorly characterized clade containing *Chalara holubovae* (Fig. 8). In addition, two closely related SHs in the genus *Rhexocercosporidium*, for which both saprotrophic and endophytic lifestyles were reported [55,56], exhibited particularly high specificity toward Poaceae (Fig. 8).

**Figure 5:**
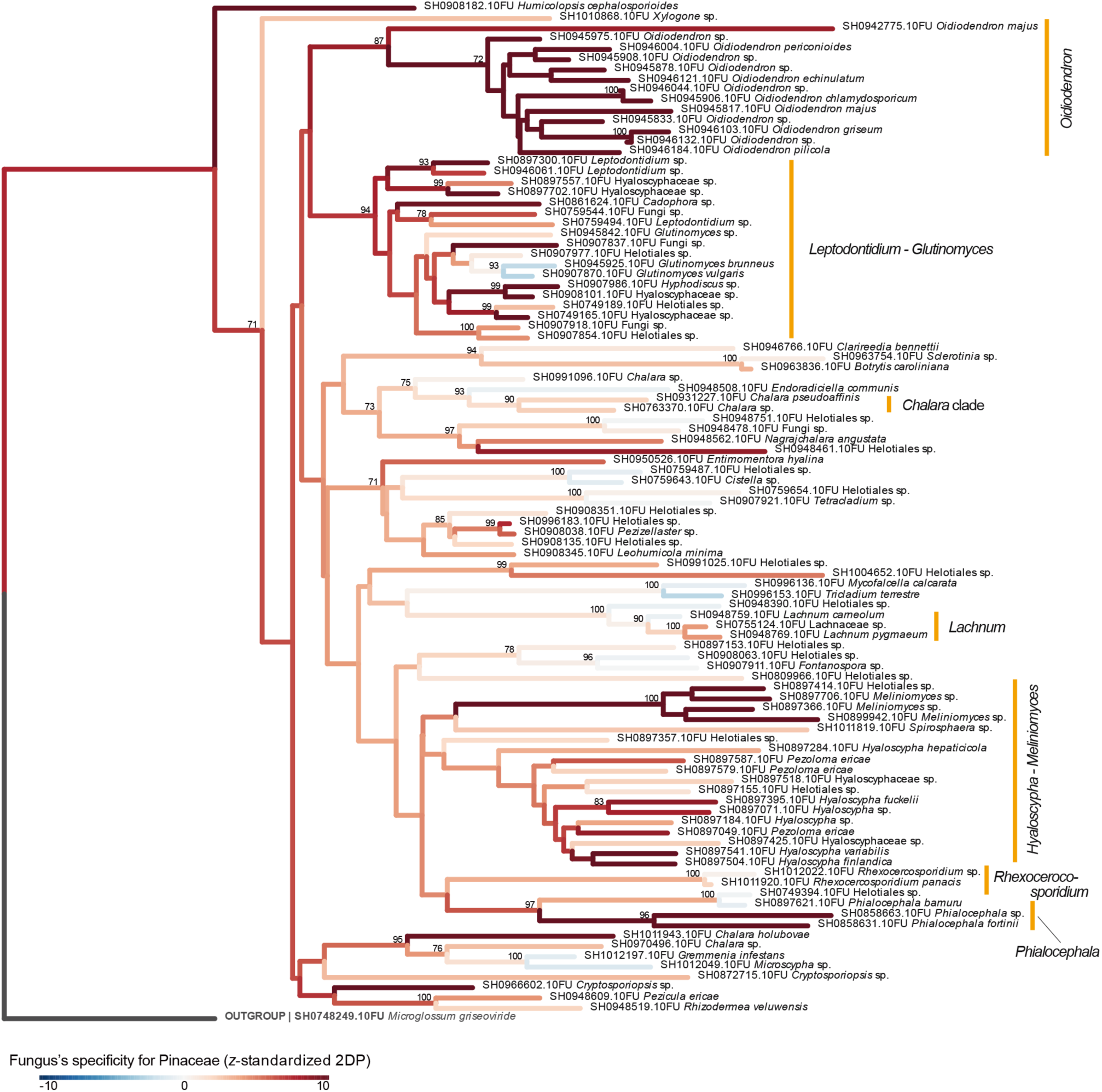
Phylogenetic patterns in the specificity of Helotiales fungi for Pinaceae. The *z*-standardized 2DP values representing the specificity of individual Helotiales SHs for Pinaceae are reflected by branch colors in the neighbor-joining phylogeny. Given that a *z*-standardized 2DP value larger than 3 or smaller than −3 roughly indicates significant specificity (Fig. S5C), branch colors are scaled between −10 and 10 to emphasize variation around the significance thresholds. Bootstrap values greater than 70% are shown. See Table S1 for the results of Pagel’s *λ* analysis of phylogenetic conservatism.

**Figure 6:**
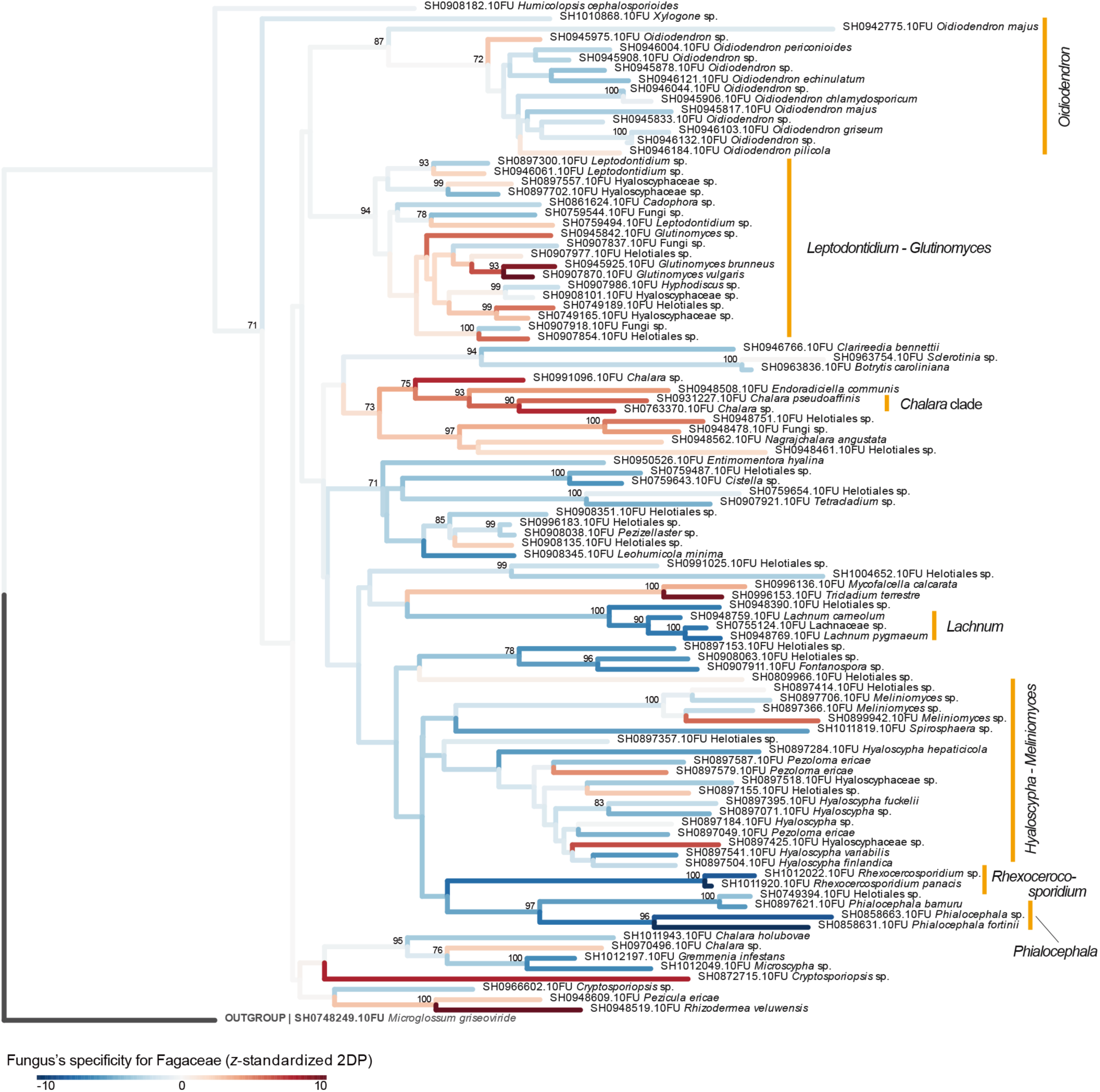
Phylogenetic patterns in the specificity of Helotiales fungi for Fagaceae. The *z*-standardized 2DP values representing the specificity of individual Helotiales SHs for Fagaceae are reflected by branch colors in the neighbor-joining phylogeny. Given that a *z*-standardized 2DP value larger than 3 or smaller than −3 roughly indicates significant specificity (Fig. S5C), branch colors are scaled between −10 and 10 to emphasize variation around the significance thresholds. Bootstrap values greater than 70% are shown. See Table S1 for the results of Pagel’s *λ* analysis of phylogenetic conservatism.

**Figure 7:**
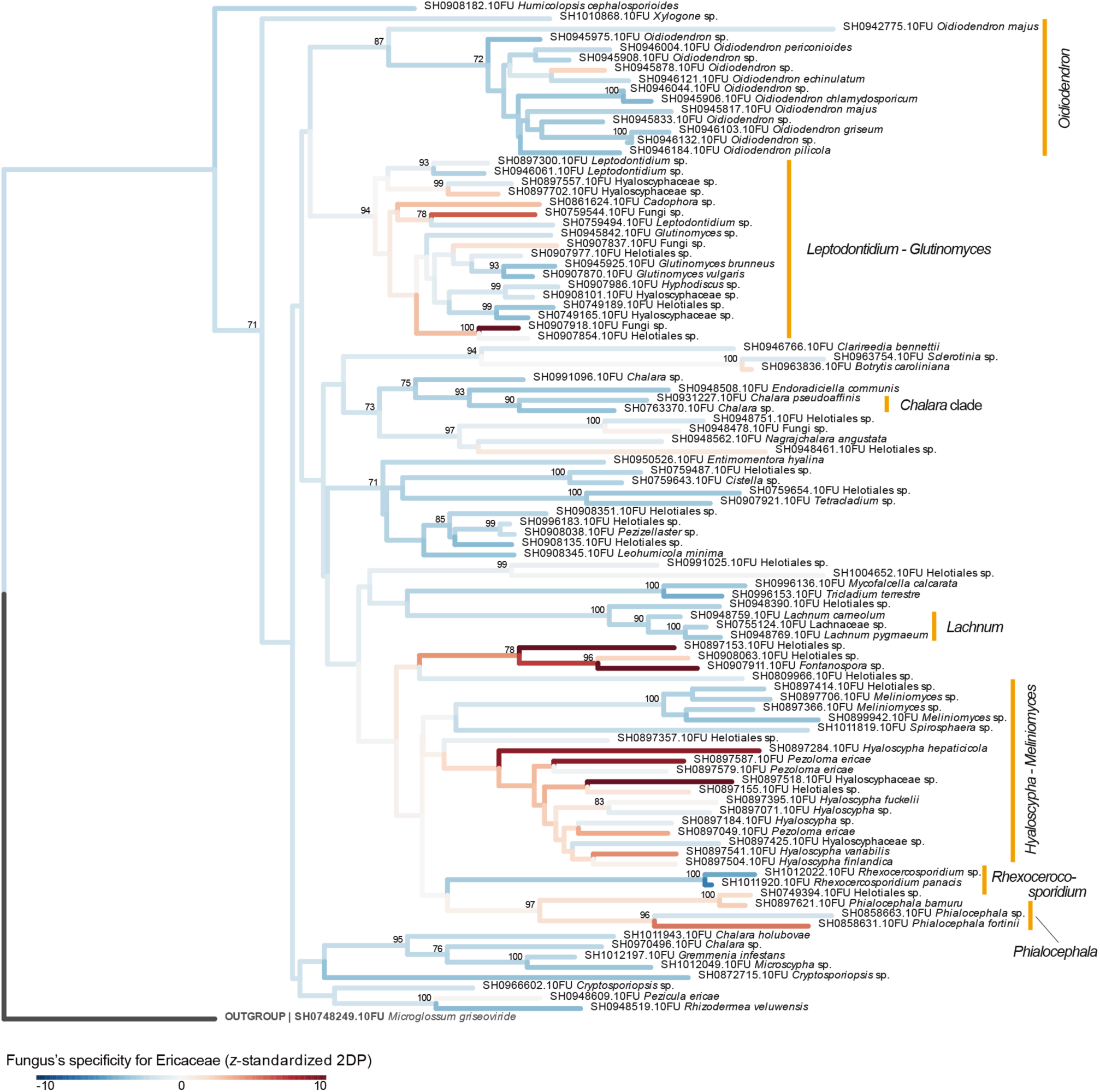
Phylogenetic patterns in the specificity of Helotiales fungi for Ericaceae. The *z*-standardized 2DP values representing the specificity of individual Helotiales SHs for Ericaceae are reflected by branch colors in the neighbor-joining phylogeny. Given that a *z*-standardized 2DP value larger than 3 or smaller than −3 roughly indicates significant specificity (Fig. S5C), branch colors are scaled between −10 and 10 to emphasize variation around the significance thresholds. Bootstrap values greater than 70% are shown. See Table S1 for the results of Pagel’s *λ* analysis of phylogenetic conservatism.

**Figure 8:**
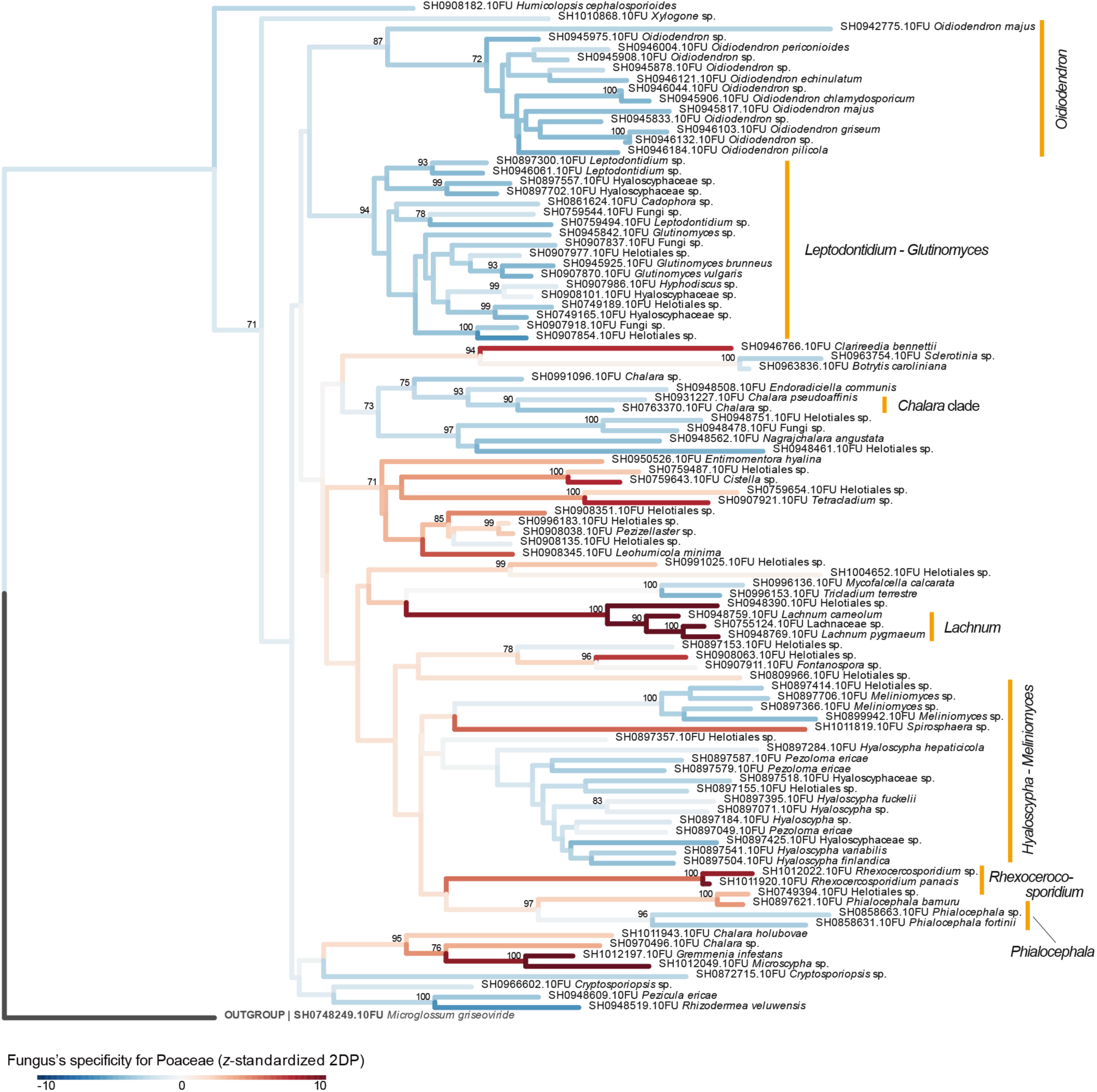
Phylogenetic patterns in the specificity of Helotiales fungi for Poaceae. The *z*-standardized 2DP values representing the specificity of individual Helotiales SHs for Poaceae are reflected by branch colors in the neighbor-joining phylogeny. Given that a *z*-standardized 2DP value larger than 3 or smaller than −3 roughly indicates significant specificity (Fig. S5C), branch colors are scaled between −10 and 10 to emphasize variation around the significance thresholds. Bootstrap values greater than 70% are shown. See Table S1 for the results of Pagel’s *λ* analysis of phylogenetic conservatism.

## DISCUSSION

By compiling a worldwide dataset of root-associated fungi, we systematically evaluated the geographic distributions and plant associations of the ecologically diverse fungal order Helotiales. Screening of the GlobalFungi database indicated that Helotiales records accompanied by associated-plant metadata remain scarce in tropical regions (Fig. 1; Fig. S1). Nonetheless, the database included Helotiales–plant association records from several pioneering studies conducted in Southeast Asia and tropical America [40,57,58], suggesting that Helotiales fungi are ubiquitous components of plant-associated mycobiomes across climatic zones ranging from boreal to tropical regions. Despite the global prevalence of Helotiales, individual lineages within the order showed distinct geographic distributions, as exemplified by the overrepresentation of *Hyaloscypha* and its synonym *Meliniomyces* in Europe and that of *Glutinomyces* in Asia (Fig. 2). Although some individual SHs showed nearly worldwide distributions (e.g., SH0759652; Fig. 1B), such background spatial patterns must be taken into account when quantifying the plant associations of Helotiales fungi.

In a statistical analysis controlling for background biogeographic patterns, we found that Helotiales genera varied greatly in their specificity for associated plant lineages (Fig. 3). Among the most frequently reported Helotiales genera, *Hyaloscypha* (synonym: *Meliniomyces*) and *Oidiodendron*, both of which include ericoid mycorrhizal and root-endophytic fungi [25,59], showed the strongest associations with Pinales. *Phialocephala*, another genus containing well-characterized dark septate endophytes [9,28], also exhibited a highly significant association with Pinales. An analysis at finer taxonomic resolution (associations between fungal SHs and plant families), combined with phylogenetic analysis, further indicated that SHs within these fungal taxa were consistently associated with Pinaceae (Fig. 5). The phylogenetic analysis also showed that high association specificity for Pinaceae was conserved in the examined Helotiales phylogeny, with some terminal lineages such as *Glutinomyces*, *Cistella*, and *Tricladium* showing exceptional negative associations with the plant family. These strong relationships, together with experimental evidence for the beneficial effects of some of these fungi on members of Pinales [35,60,61], raise the possibility that these plant–fungus associations have a deep evolutionary history, potentially dating back to the early evolution of Pinaceae in the Jurassic [62,63].

In contrast to the largely conserved patterns observed for associations with Pinaceae, associations between Helotiales fungi and other ectomycorrhizal plant families exhibited more complex phylogenetic patterns (Figs. 6; Figs. S8–10). In the analysis targeting Fagaceae, for which the largest number of Helotiales occurrence records was obtained, strong associations were confined to a few terminal lineages, such as the poorly characterized endophytic genus *Glutinomyces* [64] and the predominantly saprotrophic genus *Chalara* [65]. Similar patterns, in which high specificity was restricted to small terminal lineages, were also observed for other ectomycorrhizal plant families, including Betulaceae, Salicaceae, and Myrtaceae, although the fungal lineages showing strong associations differed among the plant families examined (Figs. S8– 10). These phylogenetically structured patterns suggest that multiple host shifts of Helotiales fungi from Pinaceae to ectomycorrhizal angiosperm lineages may have occurred during the Cretaceous and subsequent periods. This hypothetical evolutionary scenario of Helotiales–plant associations parallels the broader evolutionary history of ectomycorrhizal symbiosis, which likely originated in association with Pinaceae in the Jurassic and subsequently arose repeatedly in diverse angiosperm lineages [66,67].

The retention of substantial saprotrophic capabilities has presumably enabled Helotiales fungi to occupy unique ecological roles in terrestrial ecosystems [21,23,68,69]. Some Helotiales lineages represented by *Hyaloscypha*, for example, are considered to have established mycorrhizal symbioses with Ericaceae plants in the Early Cretaceous [67,70], potentially facilitating the diversification and persistence of Ericaceae in nutrient-poor environments in which decomposition of soil organic matter is limited [21,71]. Although ericoid mycorrhizal fungi form characteristic morphological structures (hyphal coils) within host cells [7], they can also interact with non-ericaceous plants (e.g., Pinaceae) in harsh environments such as tundra, heathlands, and solfatara fields [26,72–74]. In fact, closely related, and in some cases genetically identical, fungi now classified within *Hyaloscypha* have been experimentally shown to form ericoid mycorrhizal associations with Ericaceae and ectomycorrhizal or dark septate endophytic associations with Pinaceae [25,26,75]. Such multifunctionality and tight interactions [25,76] may underlie the counterintuitive statistical results, in which positive associations with Ericaceae were not detected for some Helotiales genera considered ericoid mycorrhizal (e.g., *Oidiodendron* and *Meliniomyces*; Fig. 3). Specificity toward ericoid mycorrhizal plants may vary within each ericoid mycorrhizal lineage (see patterns within the *Hyaloscypha* lineage in Fig. 7) [16,73], while also being influenced by specificity toward background soil environmental conditions [74].

From a broader perspective, associations with Helotiales fungi are not limited to ectomycorrhizal and ericoid mycorrhizal woody plants, but they are widespread across diverse herbaceous plants conventionally categorized as arbuscular mycorrhizal or nonmycorrhizal lineages [15,41,77]. In heathlands and alpine/arctic tundra, Poaceae plants have often been reported to interact not only with arbuscular mycorrhizal fungi but also with Helotiales dark septate endophytes [41], which may provide physiological functions complementary to those of mycorrhizal fungi. Likewise, in severely phosphorus-limited soils in alpine habitats, Helotiales endophytic fungi support phosphorus acquisition by nonmycorrhizal Brassicaceae plants [13,31]. In our analysis, some Helotiales clades encompassing saprotrophic, endophytic, or pathogenic lifestyles, such as *Lachnum* and *Rhexocercosporidium* [78,79], showed high association specificity for Poaceae (Fig. 8), whereas the specificity of Helotiales fungi for Brassicaceae was less evident (Fig. S29). Intriguingly, individual Helotiales fungi isolated from ectomycorrhizal or ericoid mycorrhizal plants can enhance the growth of arbuscular mycorrhizal and nonmycorrhizal plants when inoculated under laboratory conditions [41,80]. Therefore, even if Helotiales fungi preferentially associate with ectomycorrhizal and ericoid mycorrhizal plants over arbuscular mycorrhizal and nonmycorrhizal plants under natural conditions, they may function as flexible symbionts with broad potential host ranges. In this respect, it is crucial to understand how their physiological functions are regulated by host immune systems [81], which may differ substantially among plant lineages with contrasting mycorrhizal strategies.

Whereas this study provides an overview of Helotiales–plant associations in light of the hypothetical evolutionary history of this species-rich fungal order, the potential pitfalls and limitations of the present analysis need to be acknowledged. First, even though the GlobalFungi database offered an invaluable opportunity to systematically analyze plant–fungus associations [42], occurrence records with reliable associated-plant information need to be further increased for more comprehensive analyses. In particular, more records from tropical regions are needed to integrate insights across the full spectrum of ectomycorrhizal (e.g., Dipterocarpaceae [82]), arbuscular mycorrhizal, and nonmycorrhizal plants worldwide. Second, although the retrieval of DNA sequence data from the UNITE database allowed a systematic phylogenetic approach, the use of ITS sequence information provided limited bootstrap support for some major Helotiales lineages (Figs. 5–8). Extension to multigene or whole-genome-scale phylogenetic analyses [1] will enable more reliable inferences about the phylogenetic conservatism and ancestral states of Helotiales– plant associations. Third, to better understand the ecological flexibility of Helotiales fungi, the present analyses at the SH and genus levels should be complemented by strain- or isolate-level analyses [83]. Many SHs in our dataset exhibited worldwide distributions (e.g., Fig. 1B), potentially undergoing adaptation to local hosts and environmental conditions [83,84]. Integration of ecological analyses with population genetic analyses will provide a platform for understanding how plant– root-associated fungal interactions are continuously reshaped in terrestrial ecosystems.

## CONCLUSIONS

Based on a worldwide analysis of a fungal DNA sequence database, we herein provide an overview of associations between the ecologically diverse fungal order Helotiales and land plants. The integration of association-specificity statistics and phylogenetic analyses suggested the possibility that Helotiales has maintained long-lasting associations with Pinaceae, potentially dating back to the Jurassic [62,63]. In contrast, tight associations between Helotiales fungi and other plant families were largely restricted to terminal clades within the Helotiales phylogeny, suggesting the possibility of multiple host shifts from Pinaceae to other plant lineages. Given that Helotiales fungi potentially influence the assembly of mycorrhizal fungi in plant root systems [15,37,38] and that they may be utilized in forest ecosystem management and sustainable agriculture [30,35,36,85,86], uncovering their “cryptic” ecology and physiology is of substantial basic and applied scientific interest. Further comprehensive analyses of Helotiales–plant associations in terrestrial ecosystems, combined with laboratory inoculation experiments and subsequent transcriptomic analyses [21], will help uncover the dynamic nature of saprotrophic, endophytic, and mycorrhizal associations.

## Acknowledgments

ChatGPT (OpenAI) was used to assist with the development of analysis code and English-language editing. All AI-assisted outputs were reviewed and verified by the authors.

## Authors’ contributions

HT designed the research with ASR and MN. HT analyzed the data with AM. HT wrote the manuscript with ASR, MN, and AM. All authors read and approved the final manuscript.

## Availability of data and material

The fungal occurrence records data files (“GlobalFungi_5_SH_abundance_ITS1_ITS2.txt.gz” and “GlobalFungi_5_sample_metadata.txt.gz”) were obtained from the GlobalFungi database (https://globalfungi.com/). The ITS1 and ITS2 sequence files of fungal Species Hypotheses (“sh_general_release_s_04.04.2024.tgz” and “sh_general_release_dynamic_04.04.2024.SHs.tax.bz2”) were obtained from the UNITE database (https://unite.ut.ee/repository.php). The filtered dataset of Helotiales occurrence records with plant information are available from the Zenodo repository (https://doi.org/10.5281/zenodo.22104961). The computer codes are available from the Zenodo repository (https://doi.org/10.5281/zenodo.23003026).

## DECLARATIONS

### Ethics approval and consent to participate

Not applicable

### Consent for publication

Not applicable

## Abbreviations

2DP: two-dimensional preference
FDR: false discovery rate
ITS: internal transcribed spacer
ITS1: internal transcribed spacer 1
ITS2: internal transcribed spacer 2
PCWDE: plant cell wall-degrading enzyme
rRNA: ribosomal RNA
SD: standard deviation
SH: Species Hypothesis

