## Supplementary Figures and Tables for "Global distributions and plant associations of the ecologically diverse ascomycete order Helotiales"

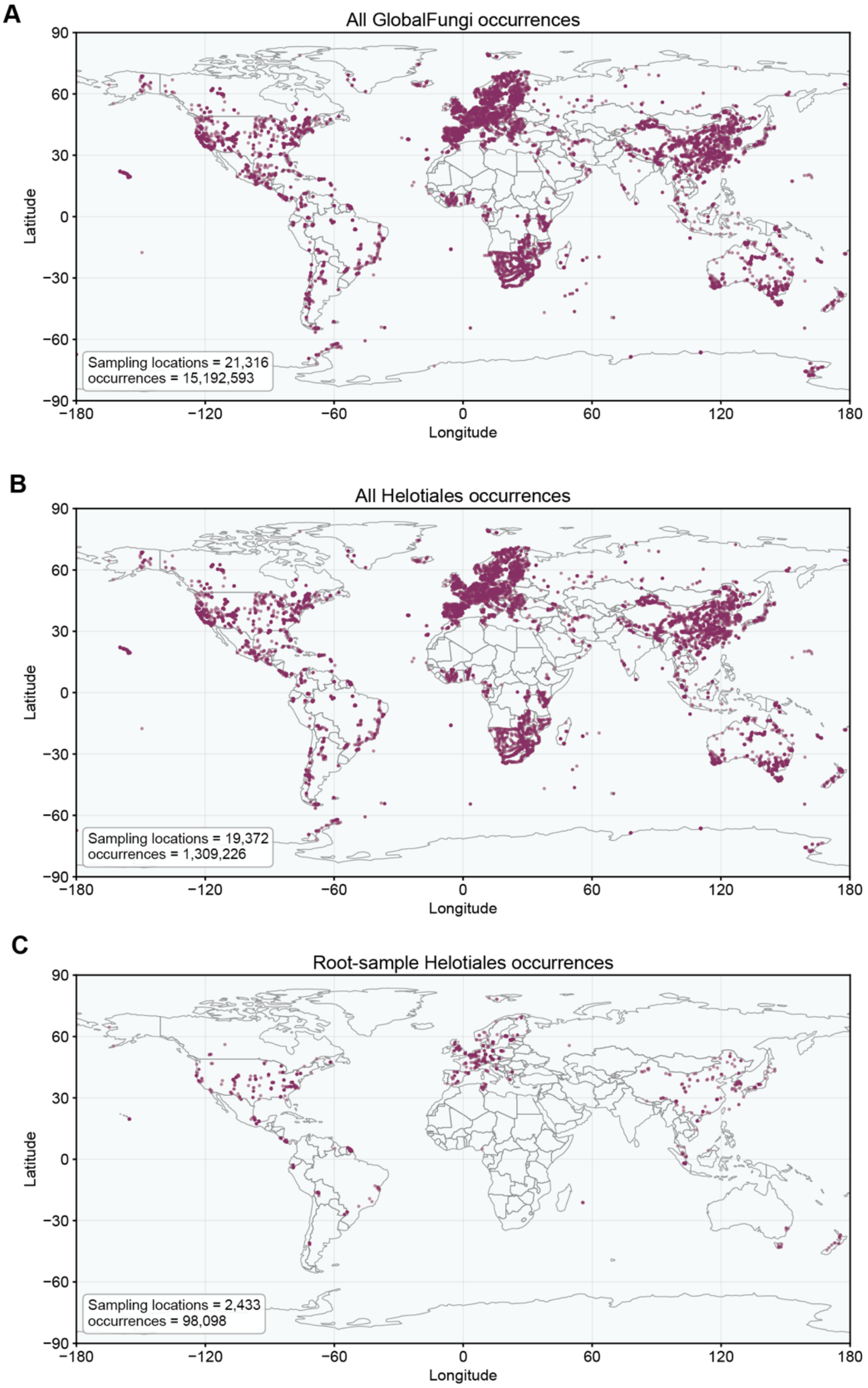


**Figure S1**: Map of occurrence records in the GlobalFungi database. (A) Map of all fungal occurrences in the GlobalFungi 5.0 database. (B) Map of all the sampling locations at which Helotiales sequences were obtained. (C) Map of all the sampling locations from which root samples with Helotiales sequences were obtained.

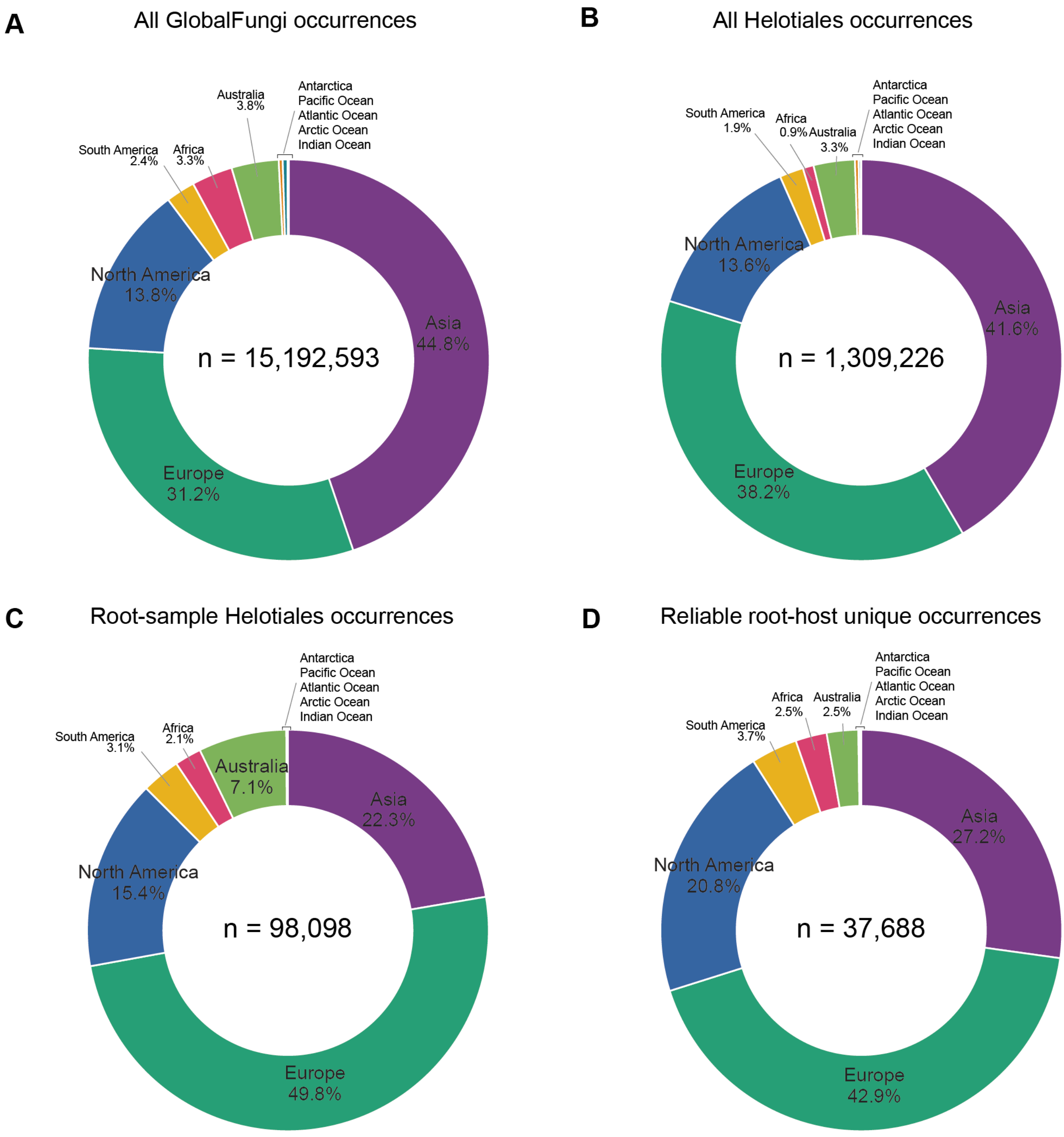


**Figure S2**: Composition of occurrence records by geographic regions. (A) All samples with fungal sequences. (B) All samples with Helotiales sequences. (C) All root samples with Helotiales sequences. (D) Unique Helotiales occurrence records with reliable host plant information.

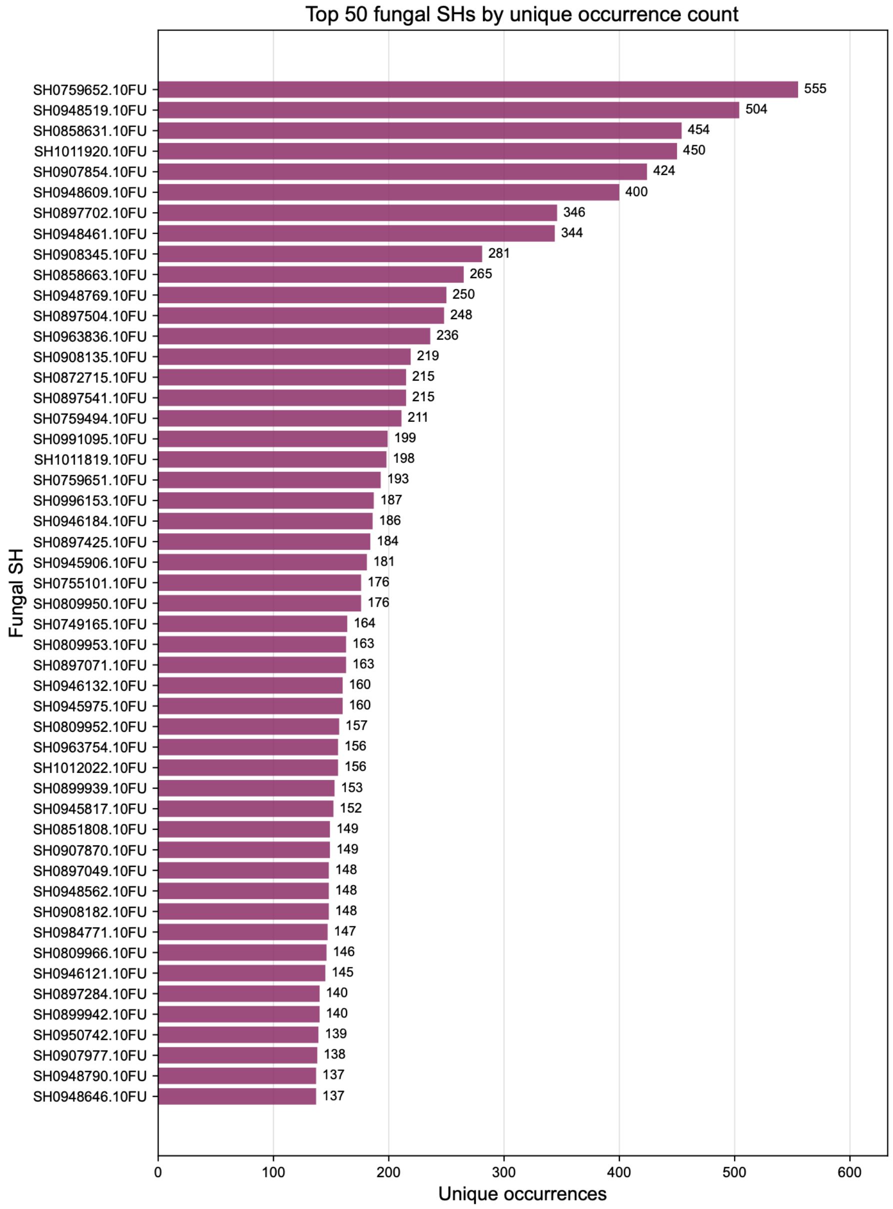


**Figure S3**: Number of unique occurrences for the 50 most frequently observed Helotiales SHs. Unique occurrence counts are shown for individual SHs.

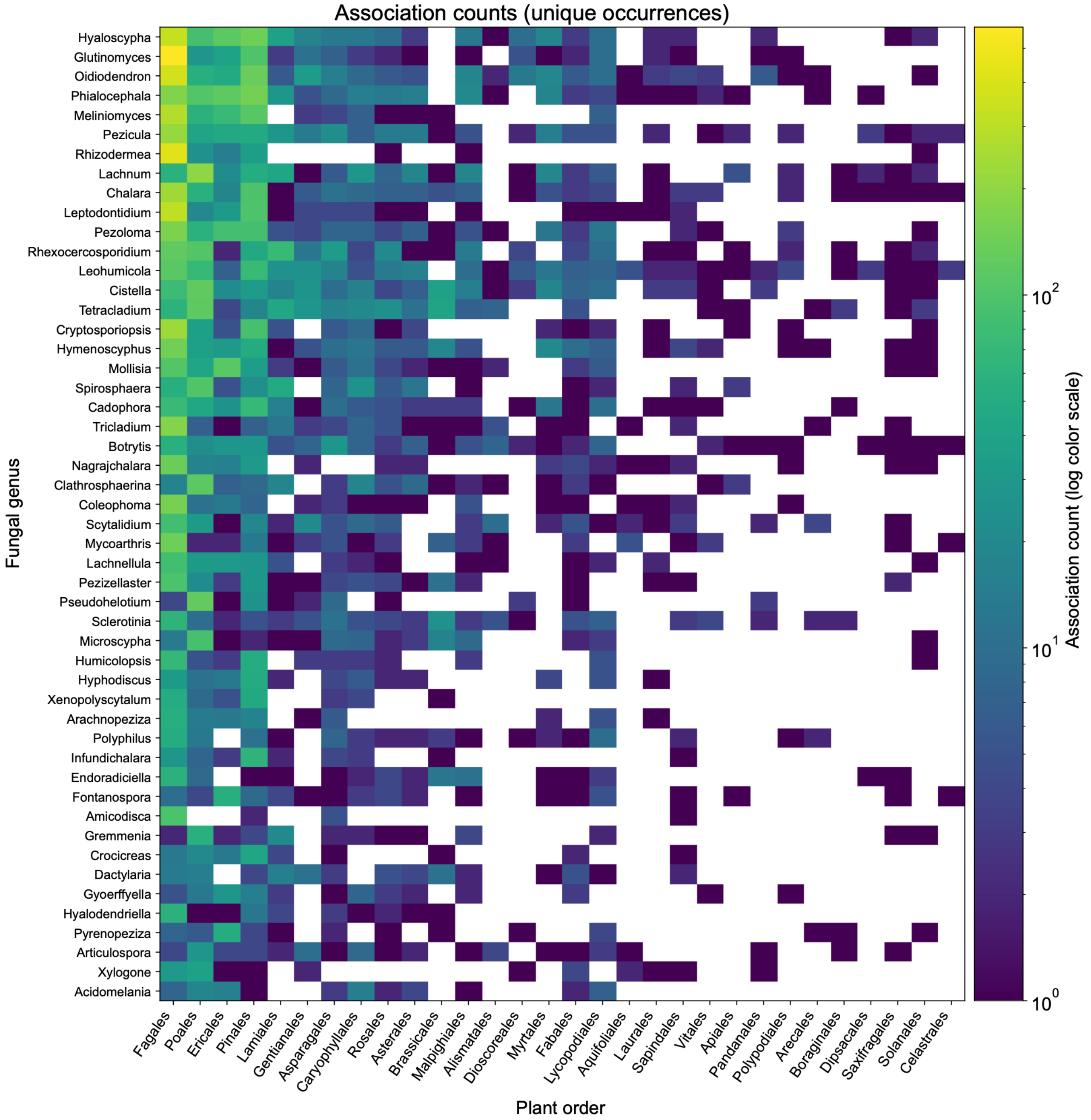


**Figure S4**: Association counts. The number of unique occurrences is shown for each combination of fungal genus and plant order.

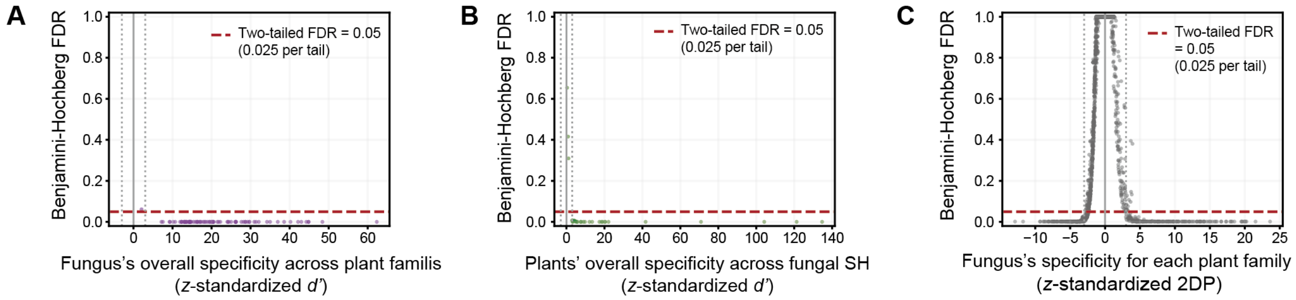


**Figure S5**: Statistical significance of association specificity between Helotiales SHs and plant families. For the analysis shown in Figure 4, the background information of the relationship between index values and false discovery rate (FDR) is shown. (A) Statistical significance of the $z$-standardized $d'$ metric for Helotiales SHs. A $z$-standardized $d'$ value larger than 3 or smaller than −3 roughly indicates significant specificity after Benjamini–Hochberg adjustment of $P$-values, with a two-tailed FDR threshold of 0.05 (0.025 per tail). (B) Statistical significance of the $z$-standardized $d'$ metric for plant families. (C) Statistical significance of the $z$-standardized 2DP metric.

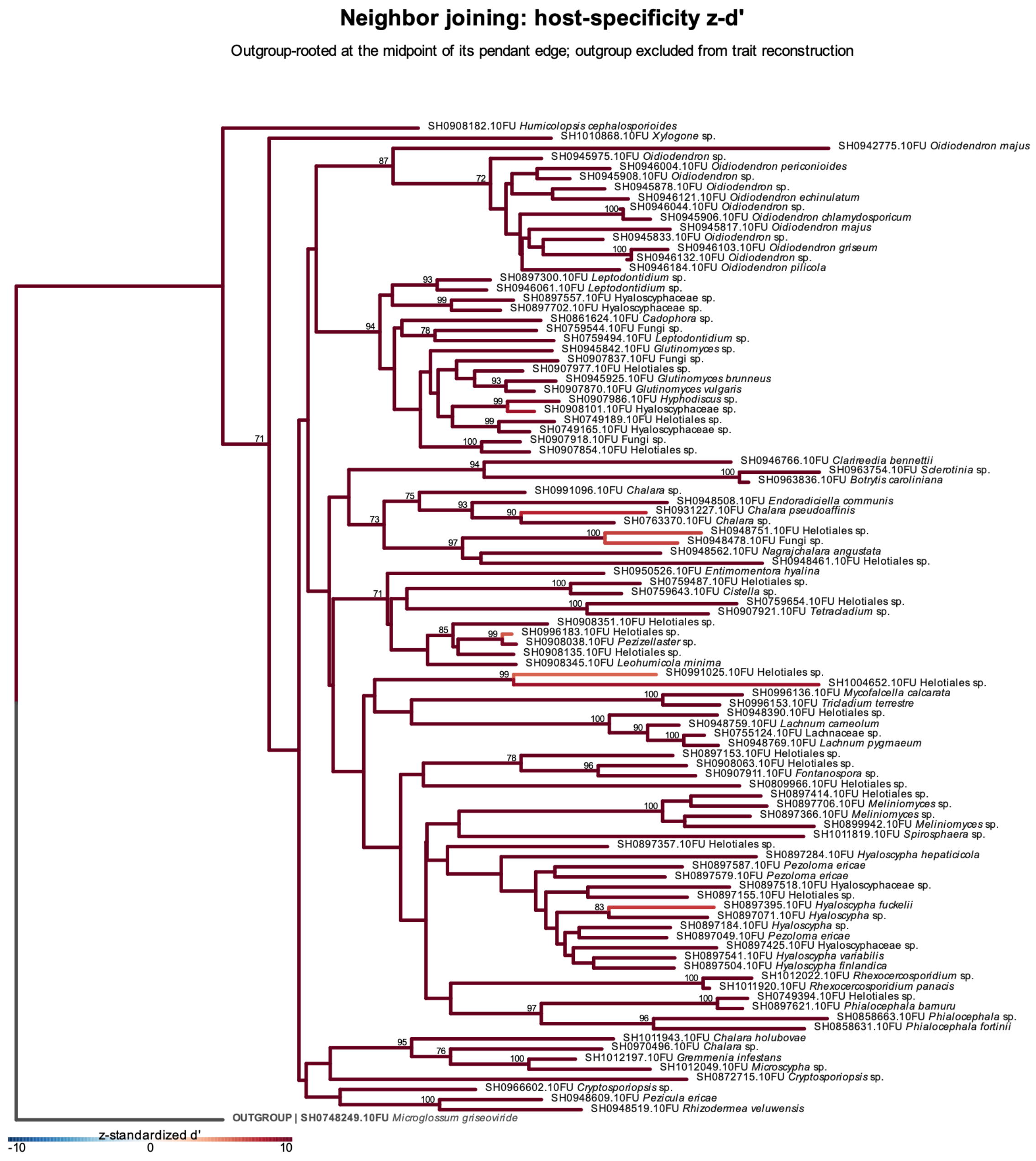


**Figure S6**: Phylogenetic patterns in the specificity of Helotiales fungi across plant families. The *z*-standardized $d'$ values representing the specificity of individual Helotiales SHs across plant families are reflected by branch colors in the neighbor-joining phylogeny. Branch colors are scaled between −10 and 10 to emphasize variation around the significance thresholds. Bootstrap values greater than 70% are shown.

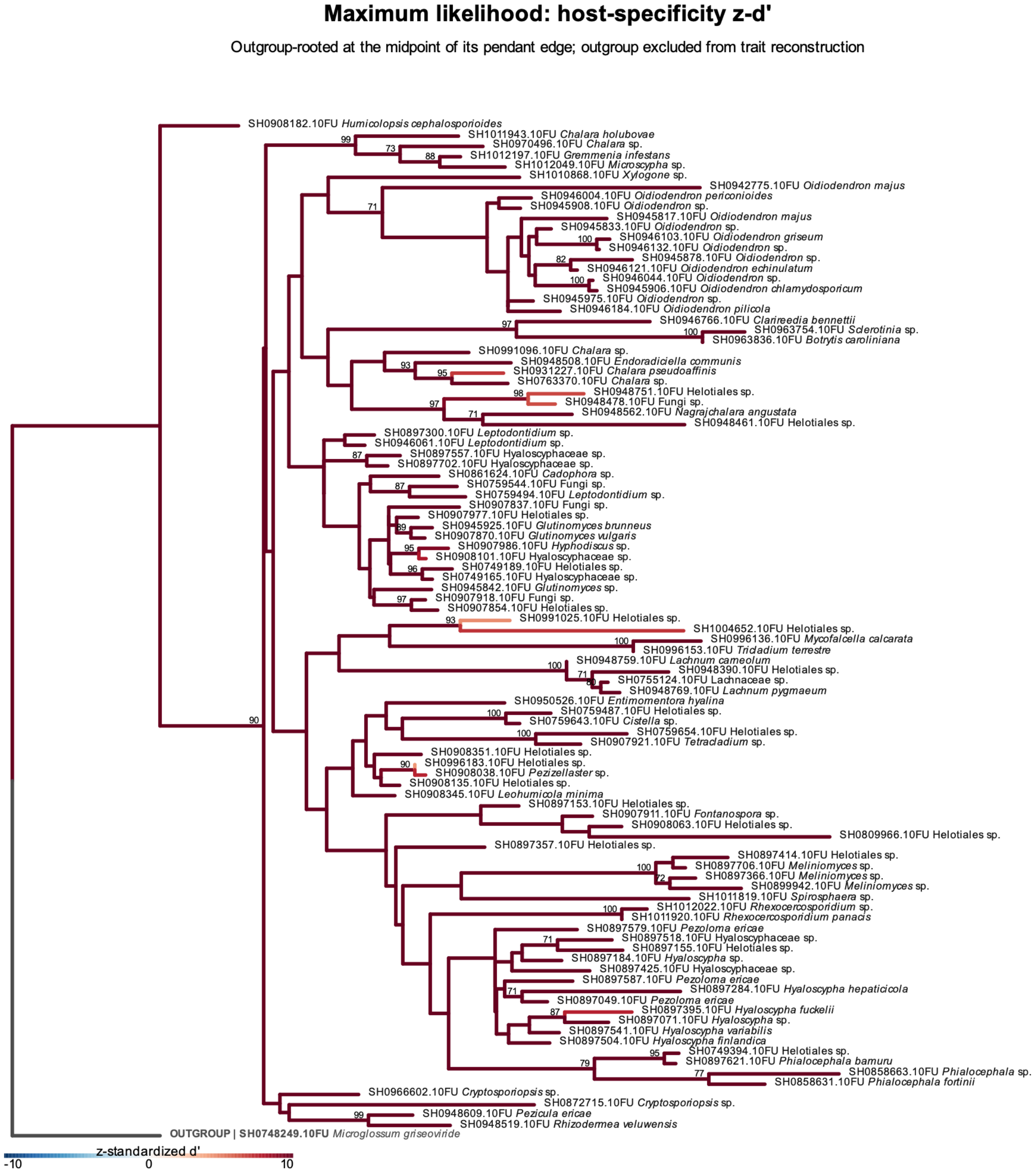


**Figure S7**: Phylogenetic patterns in the specificity of Helotiales fungi across plant families. The *z*-standardized *d′* values representing the specificity of individual Helotiales SHs across plant families are reflected by branch colors in the maximum-likelihood phylogeny. Branch colors are scaled between −10 and 10 to emphasize variation around the significance thresholds. Bootstrap values greater than 70% are shown.

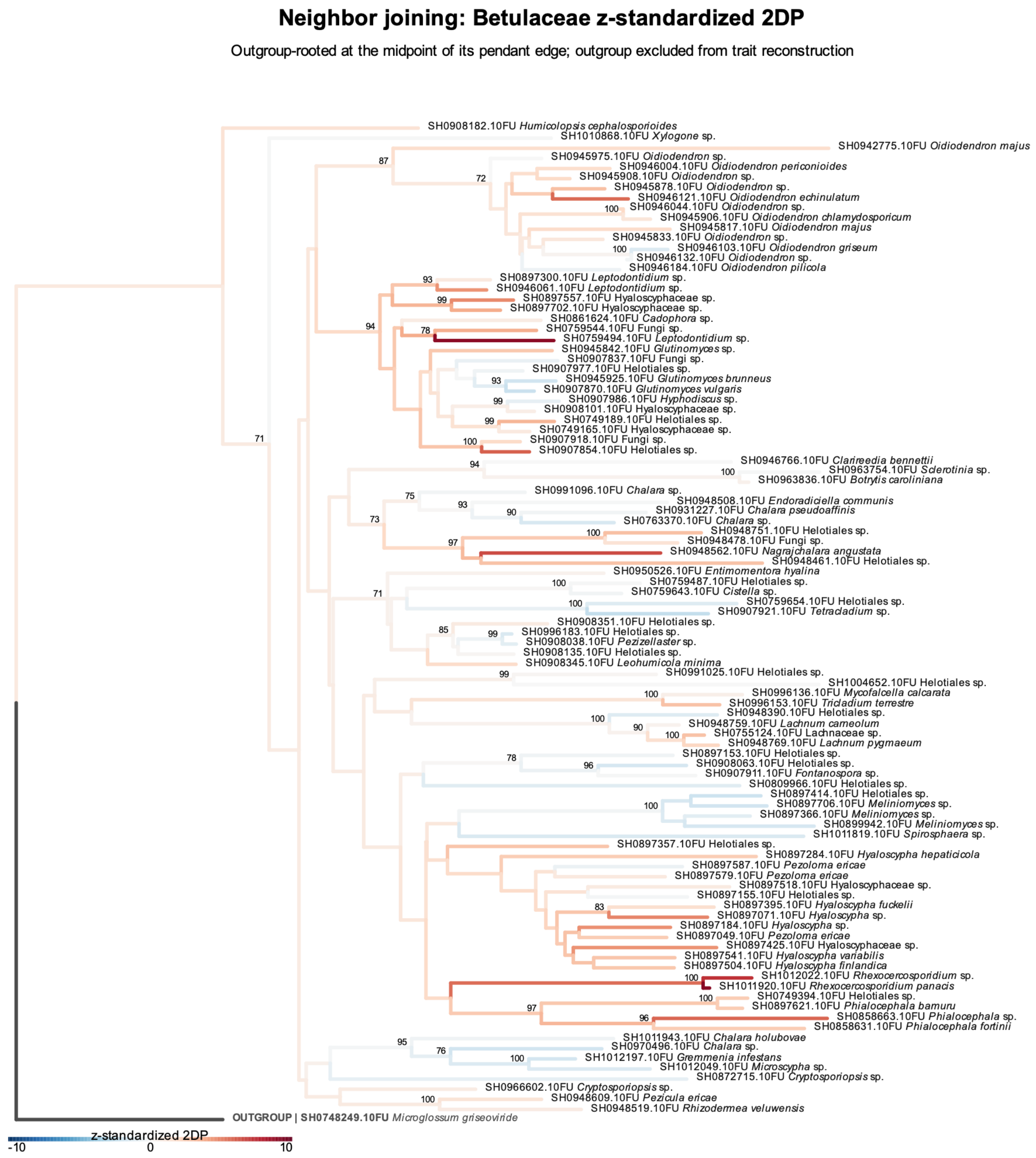


**Figure S8**: Phylogenetic patterns in the specificity of Helotiales fungi for Betulaceae. The *z*-standardized 2DP values representing the specificity of individual Helotiales SHs for Betulaceae are reflected by branch colors in the neighbor-joining phylogeny. Branch colors are scaled between −10 and 10 to emphasize variation around the significance thresholds. Bootstrap values greater than 70% are shown. See Table S1 for the results of Pagel's $\lambda$ analysis of phylogenetic conservatism.

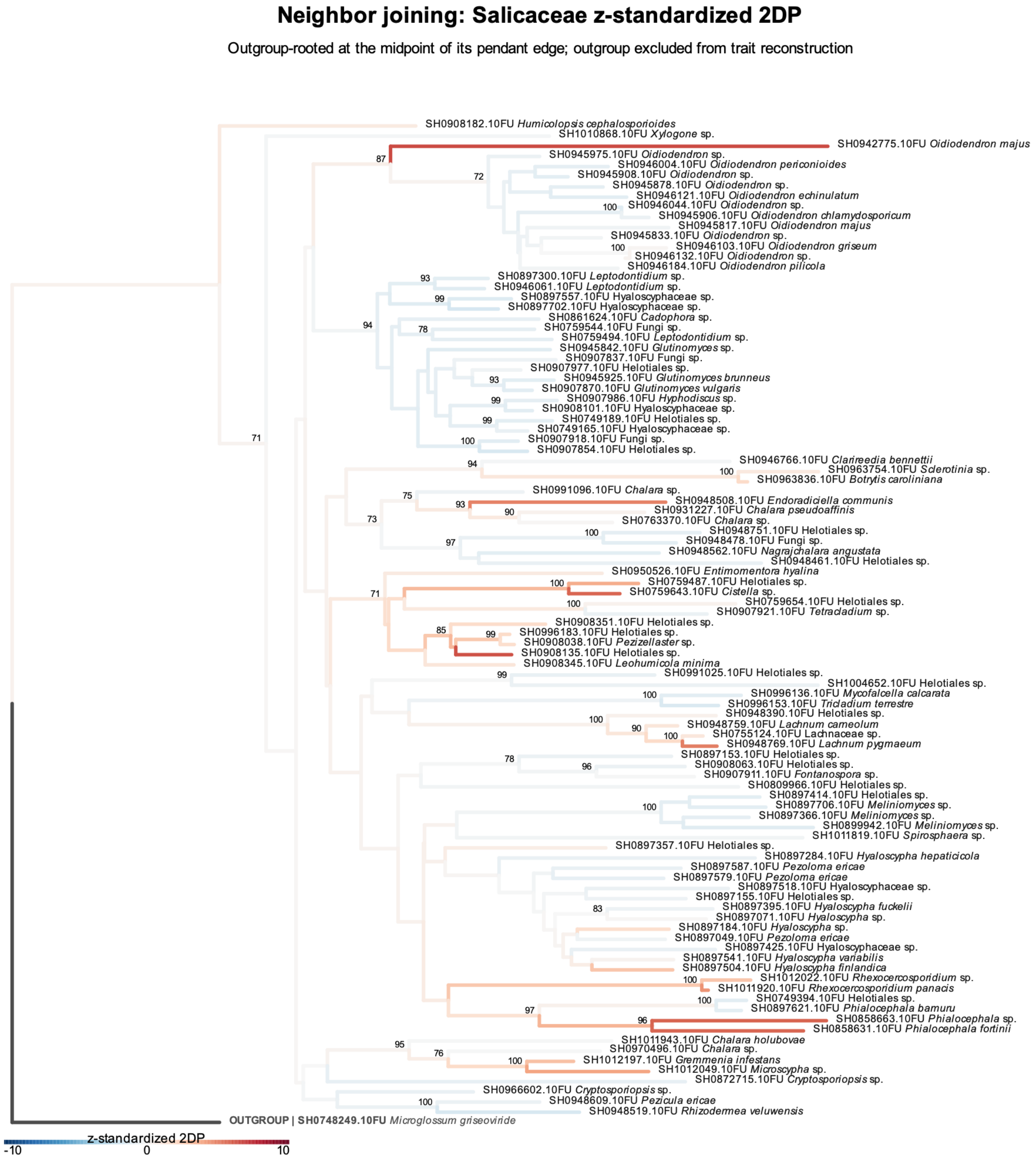


Figure S9: Phylogenetic patterns in the specificity of Helotiales fungi for Salicaceae. The *z*-standardized 2DP values representing the specificity of individual Helotiales SHs for Salicaceae are reflected by branch colors in the neighbor-joining phylogeny. Branch colors are scaled between −10 and 10 to emphasize variation around the significance thresholds. Bootstrap values greater than 70% are shown. See Table S1 for the results of Pagel's $\lambda$ analysis of phylogenetic conservatism.

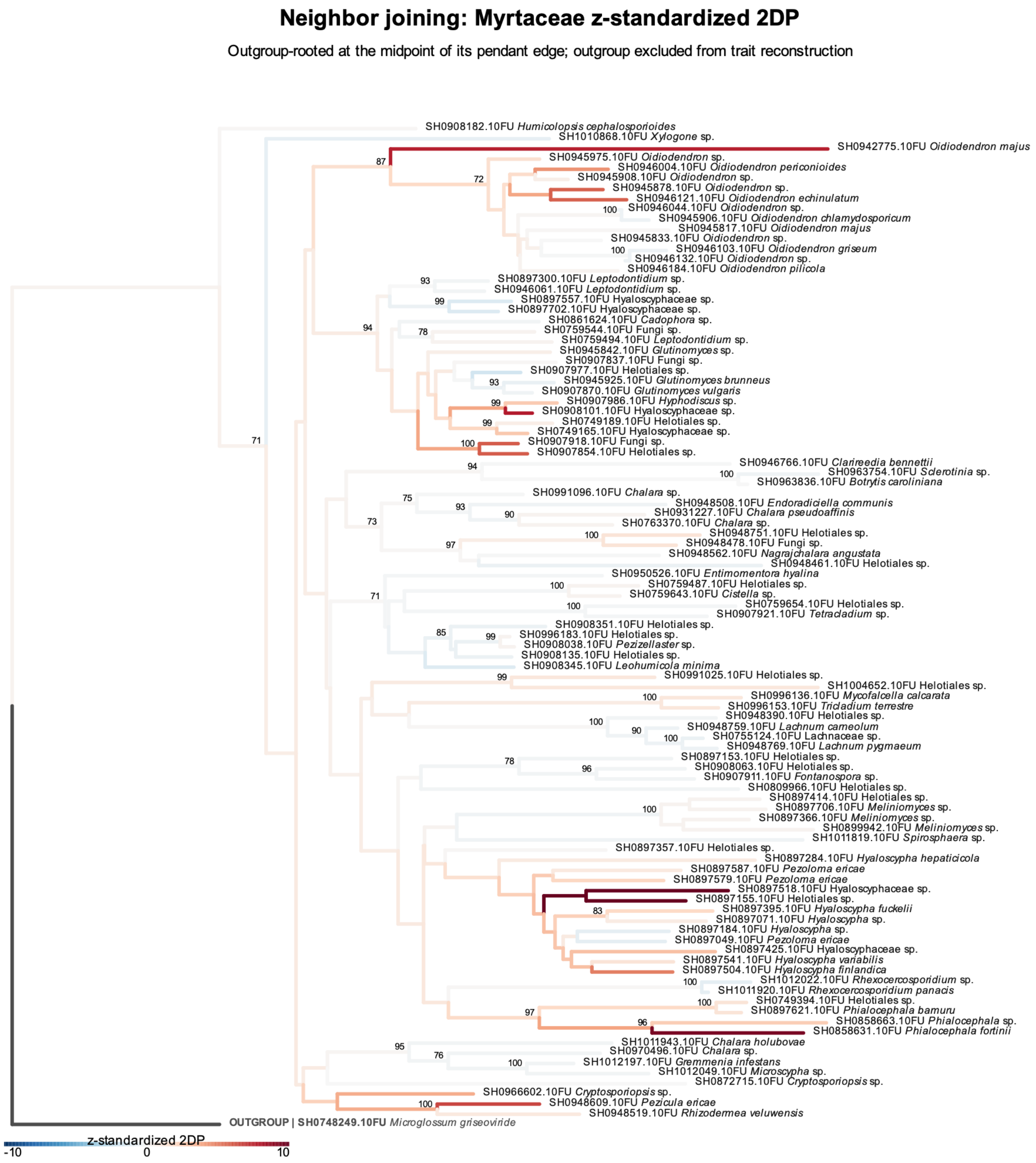


Figure S10: Phylogenetic patterns in the specificity of Helotiales fungi for Myrtaceae. The $z$-standardized 2DP values representing the specificity of individual Helotiales SHs for Myrtaceae are reflected by branch colors in the neighbor-joining phylogeny. Branch colors are scaled between −10 and 10 to emphasize variation around the significance thresholds. Bootstrap values greater than 70% are shown. See Table S1 for the results of Pagel's $\lambda$ analysis of phylogenetic conservatism.

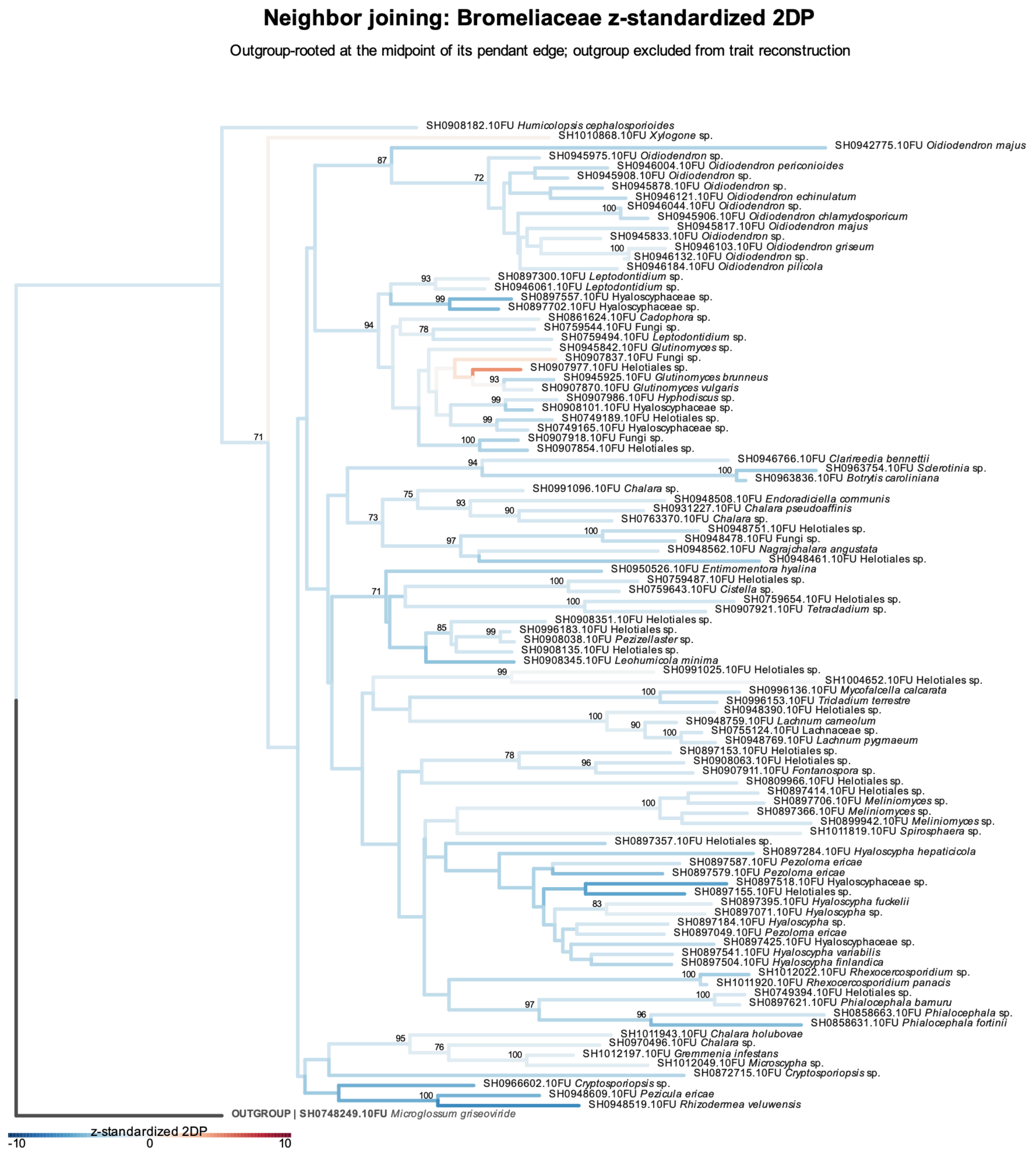


Figure S11: Phylogenetic patterns in the specificity of Helotiales fungi for Bromeliaceae. The $z$-standardized 2DP values representing the specificity of individual Helotiales SHs for Bromeliaceae are reflected by branch colors in the neighbor-joining phylogeny. Branch colors are scaled between −10 and 10 to emphasize variation around the significance thresholds. Bootstrap values greater than 70% are shown. See Table S1 for the results of Pagel's $\lambda$ analysis of phylogenetic conservatism.

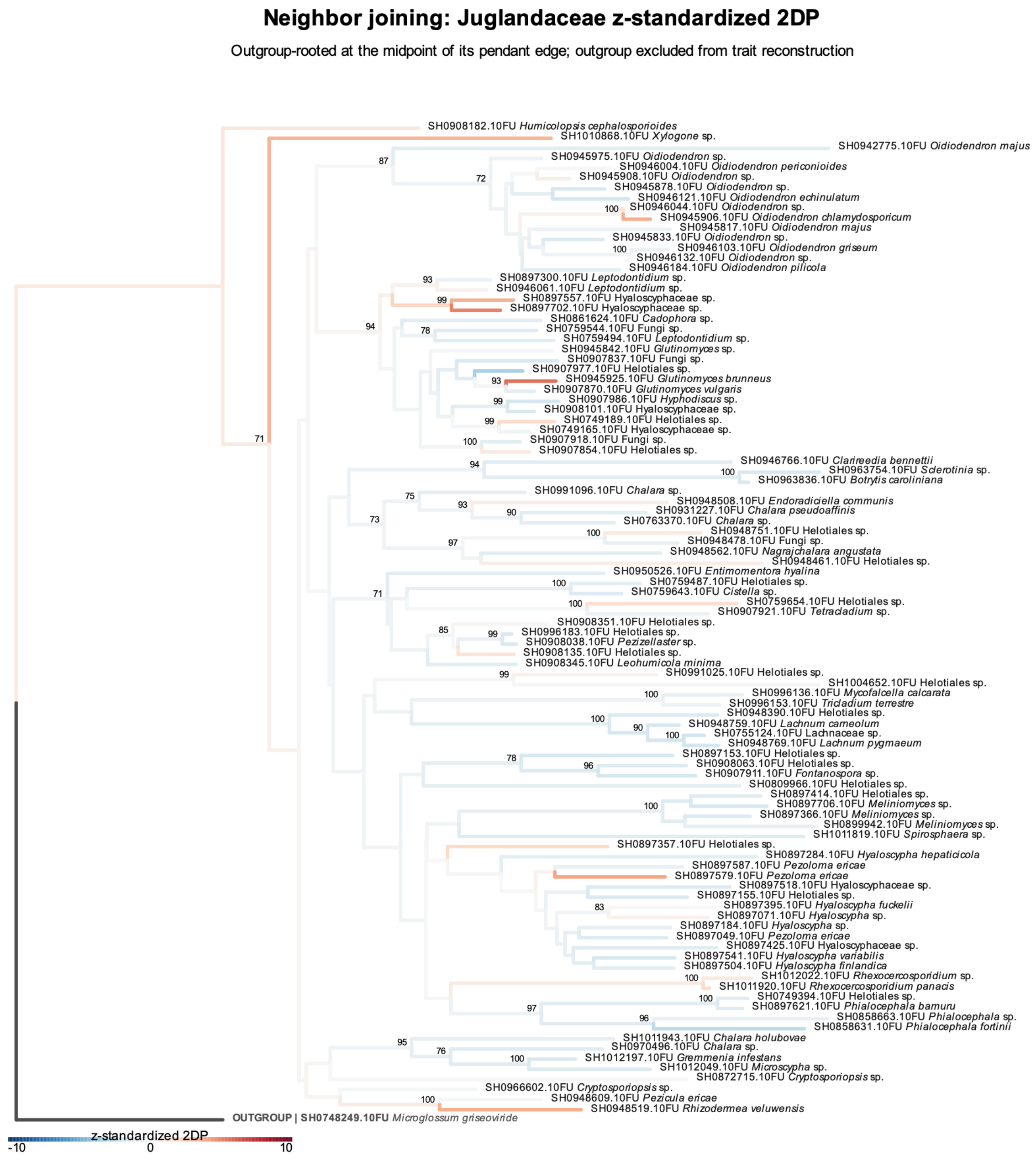


Figure S12: Phylogenetic patterns in the specificity of Helotiales fungi for Juglandaceae. The $z$-standardized 2DP values representing the specificity of individual Helotiales SHs for Juglandaceae are reflected by branch colors in the neighbor-joining phylogeny. Branch colors are scaled between −10 and 10 to emphasize variation around the significance thresholds. Bootstrap values greater than 70% are shown. See Table S1 for the results of Pagel's $\lambda$ analysis of phylogenetic conservatism.

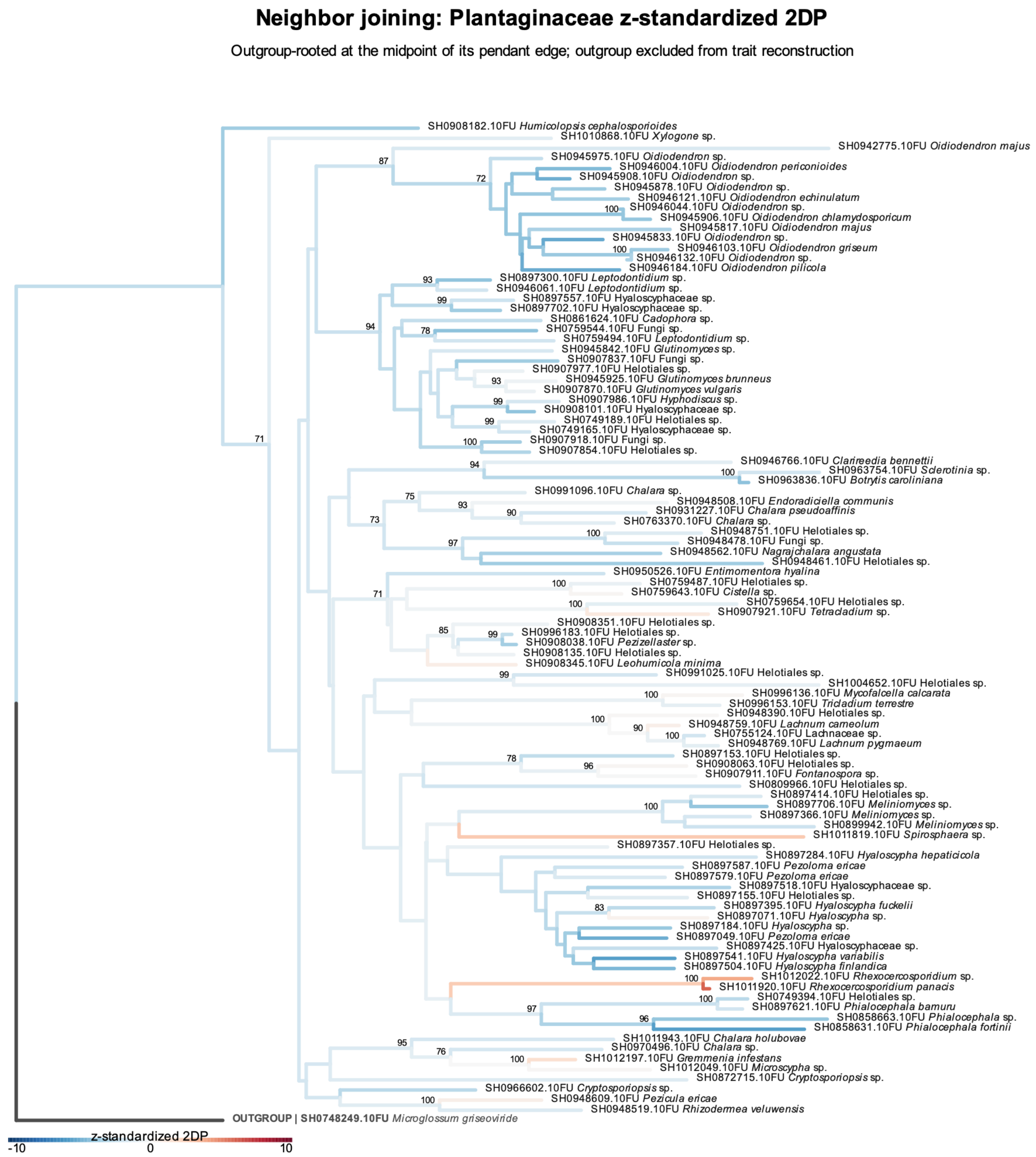


Figure S13: Phylogenetic patterns in the specificity of Helotiales fungi for Plantaginaceae. The *z*-standardized 2DP values representing the specificity of individual Helotiales SHs for Plantaginaceae are reflected by branch colors in the neighbor-joining phylogeny. Branch colors are scaled between −10 and 10 to emphasize variation around the significance thresholds. Bootstrap values greater than 70% are shown. See Table S1 for the results of Pagel's $\lambda$ analysis of phylogenetic conservatism.

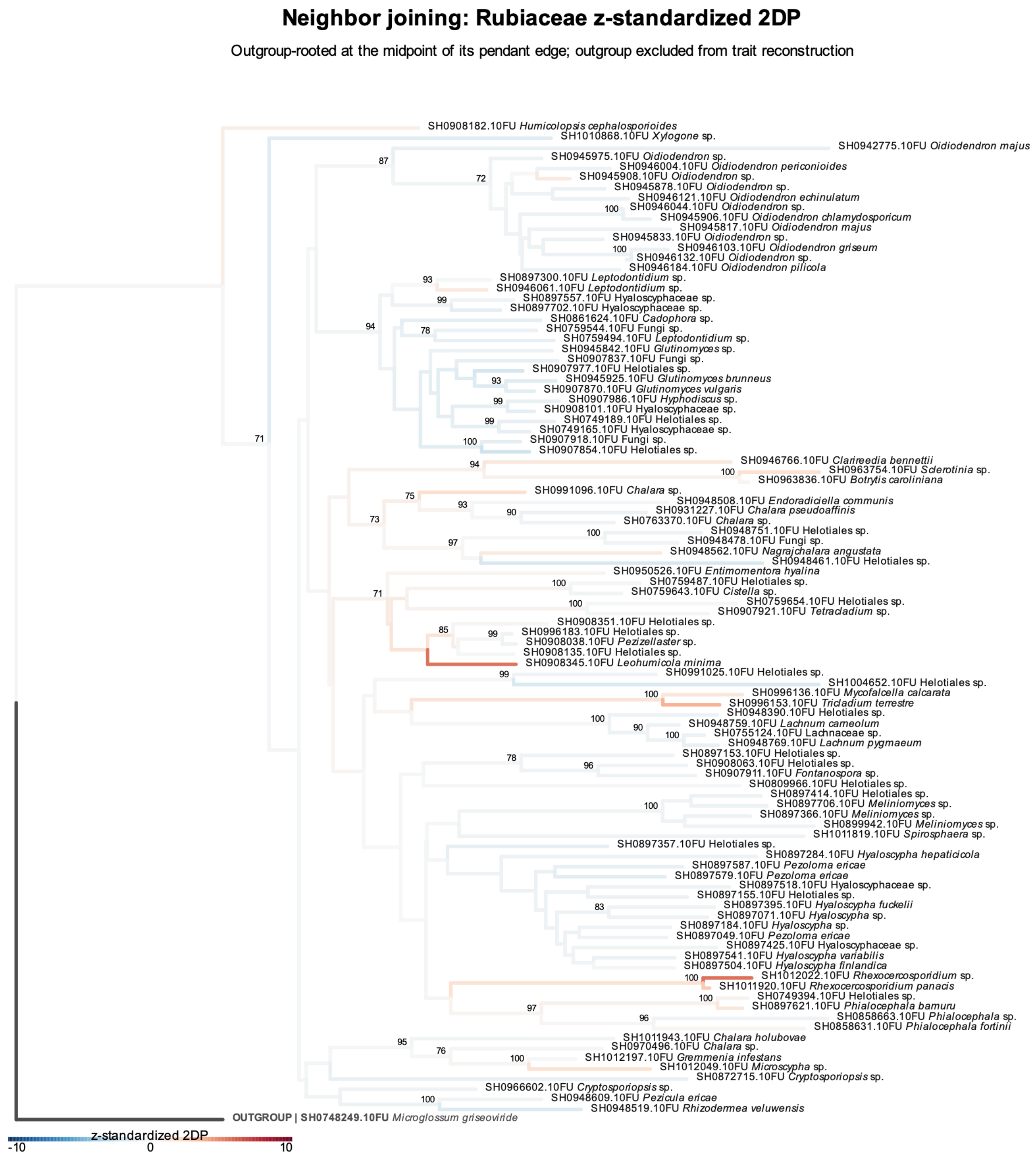


Figure S14: Phylogenetic patterns in the specificity of Helotiales fungi for Rubiaceae. The *z*-standardized 2DP values representing the specificity of individual Helotiales SHs for Rubiaceae are reflected by branch colors in the neighbor-joining phylogeny. Branch colors are scaled between −10 and 10 to emphasize variation around the significance thresholds. Bootstrap values greater than 70% are shown. See Table S1 for the results of Pagel's $\lambda$ analysis of phylogenetic conservatism.

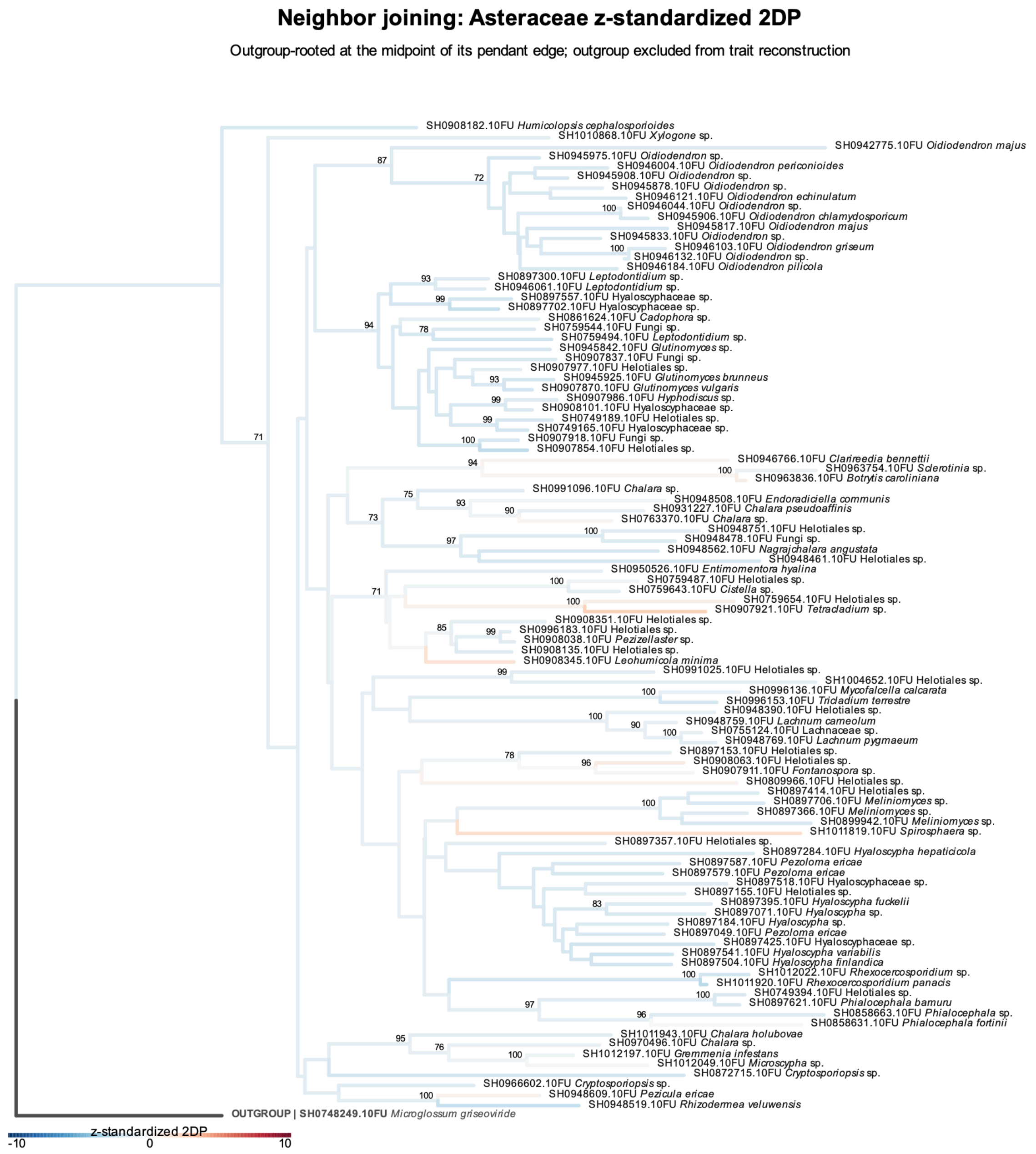


Figure S15: Phylogenetic patterns in the specificity of Helotiales fungi for Asteraceae. The *z*-standardized 2DP values representing the specificity of individual Helotiales SHs for Asteraceae are reflected by branch colors in the neighbor-joining phylogeny. Branch colors are scaled between −10 and 10 to emphasize variation around the significance thresholds. Bootstrap values greater than 70% are shown. See Table S1 for the results of Pagel's $\lambda$ analysis of phylogenetic conservatism.

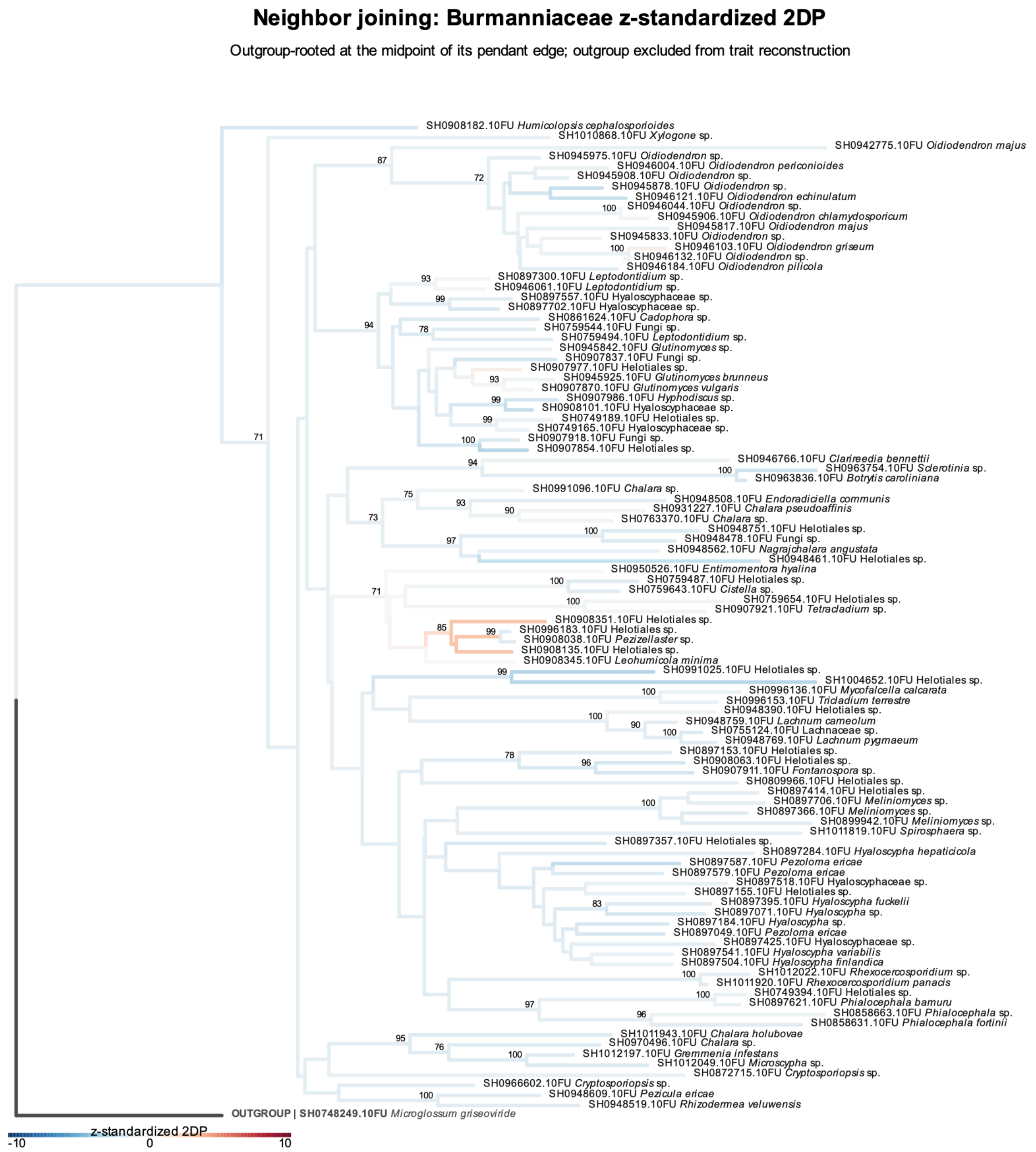


Figure S16: Phylogenetic patterns in the specificity of Helotiales fungi for Burmanniaceae. The $z$-standardized 2DP values representing the specificity of individual Helotiales SHs for Burmanniaceae are reflected by branch colors in the neighbor-joining phylogeny. Branch colors are scaled between −10 and 10 to emphasize variation around the significance thresholds. Bootstrap values greater than 70% are shown. See Table S1 for the results of Pagel's $\lambda$ analysis of phylogenetic conservatism.

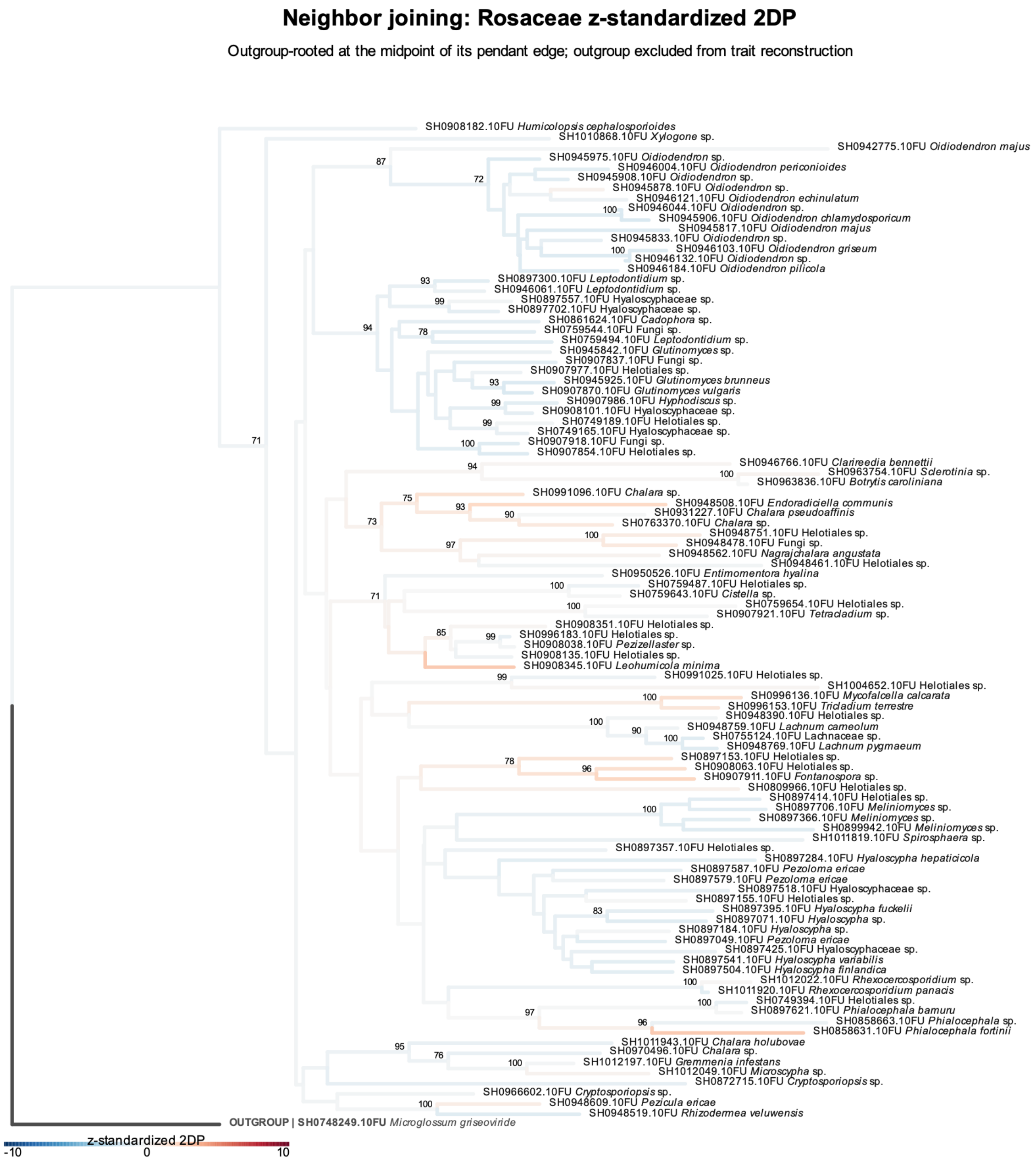


Figure S17: Phylogenetic patterns in the specificity of Helotiales fungi for Rosaceae. The $z$-standardized 2DP values representing the specificity of individual Helotiales SHs for Rosaceae are reflected by branch colors in the neighbor-joining phylogeny. Branch colors are scaled between −10 and 10 to emphasize variation around the significance thresholds. Bootstrap values greater than 70% are shown. See Table S1 for the results of Pagel's $\lambda$ analysis of phylogenetic conservatism.

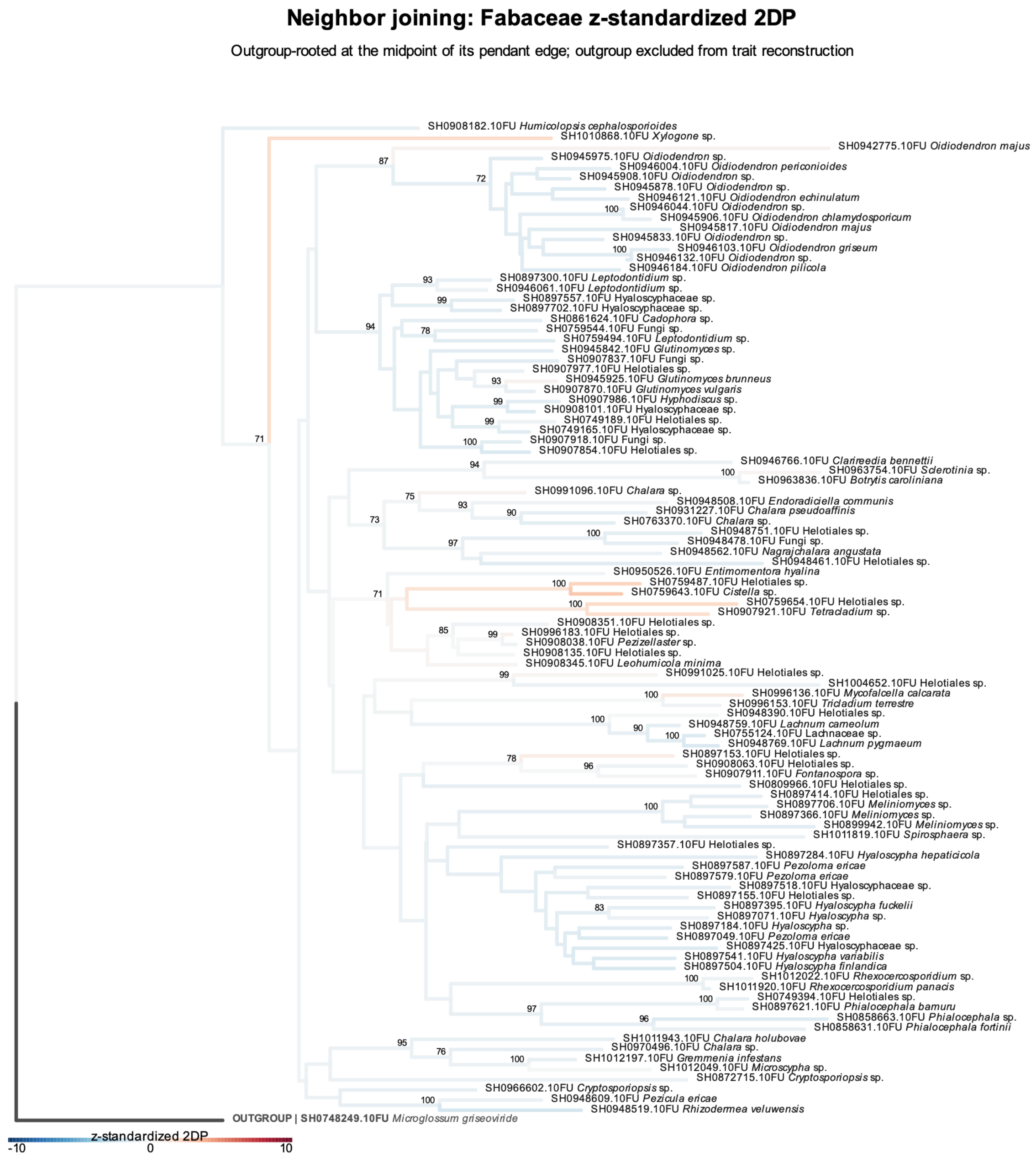


Figure S18: Phylogenetic patterns in the specificity of Helotiales fungi for Fabaceae. The *z*-standardized 2DP values representing the specificity of individual Helotiales SHs for Fabaceae are reflected by branch colors in the neighbor-joining phylogeny. Branch colors are scaled between −10 and 10 to emphasize variation around the significance thresholds. Bootstrap values greater than 70% are shown. See Table S1 for the results of Pagel's $\lambda$ analysis of phylogenetic conservatism.

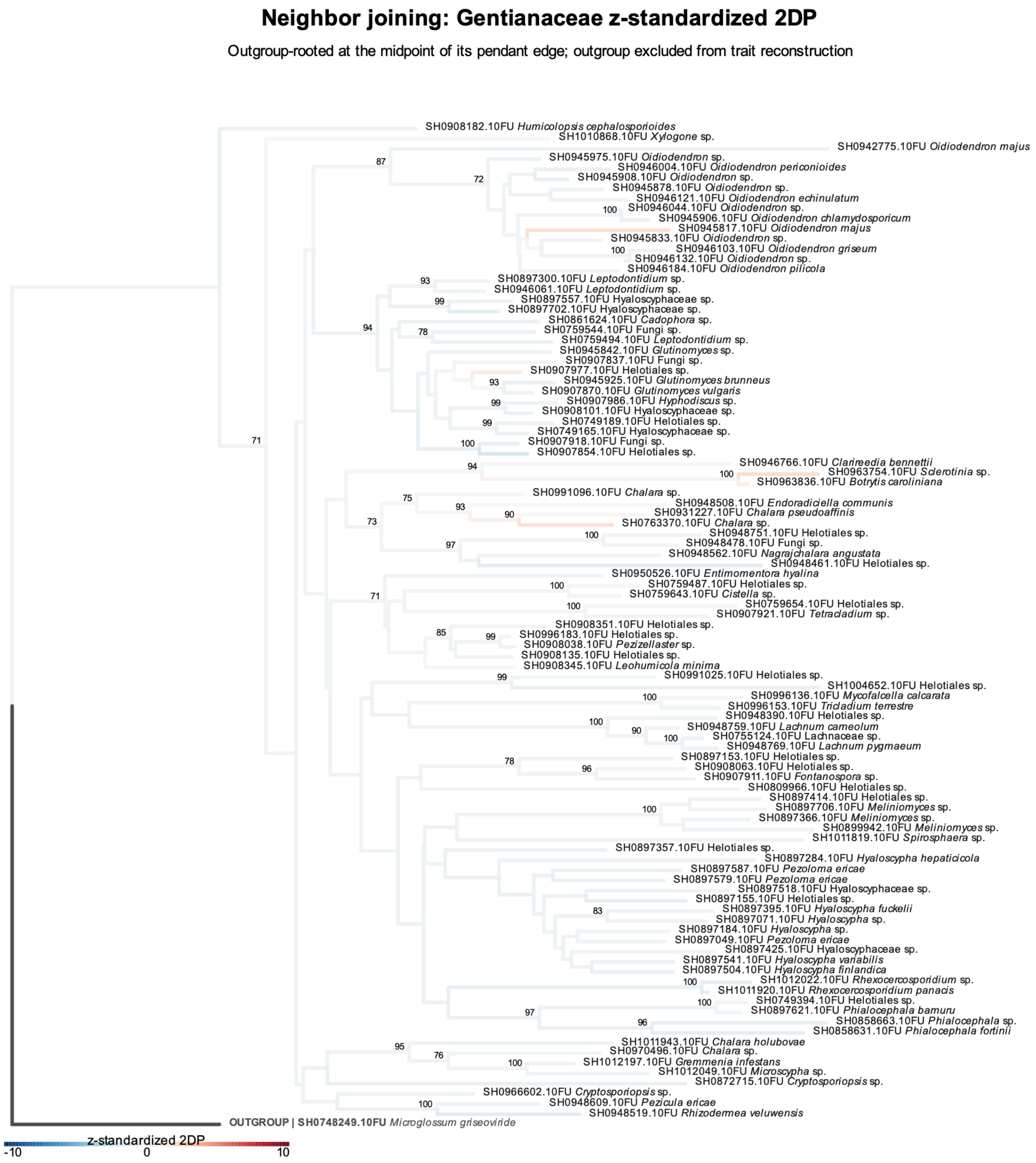


Figure S19: Phylogenetic patterns in the specificity of Helotiales fungi for Gentianaceae. The $z$-standardized 2DP values representing the specificity of individual Helotiales SHs for Gentianaceae are reflected by branch colors in the neighbor-joining phylogeny. Branch colors are scaled between −10 and 10 to emphasize variation around the significance thresholds. Bootstrap values greater than 70% are shown. See Table S1 for the results of Pagel’s $\lambda$ analysis of phylogenetic conservatism.

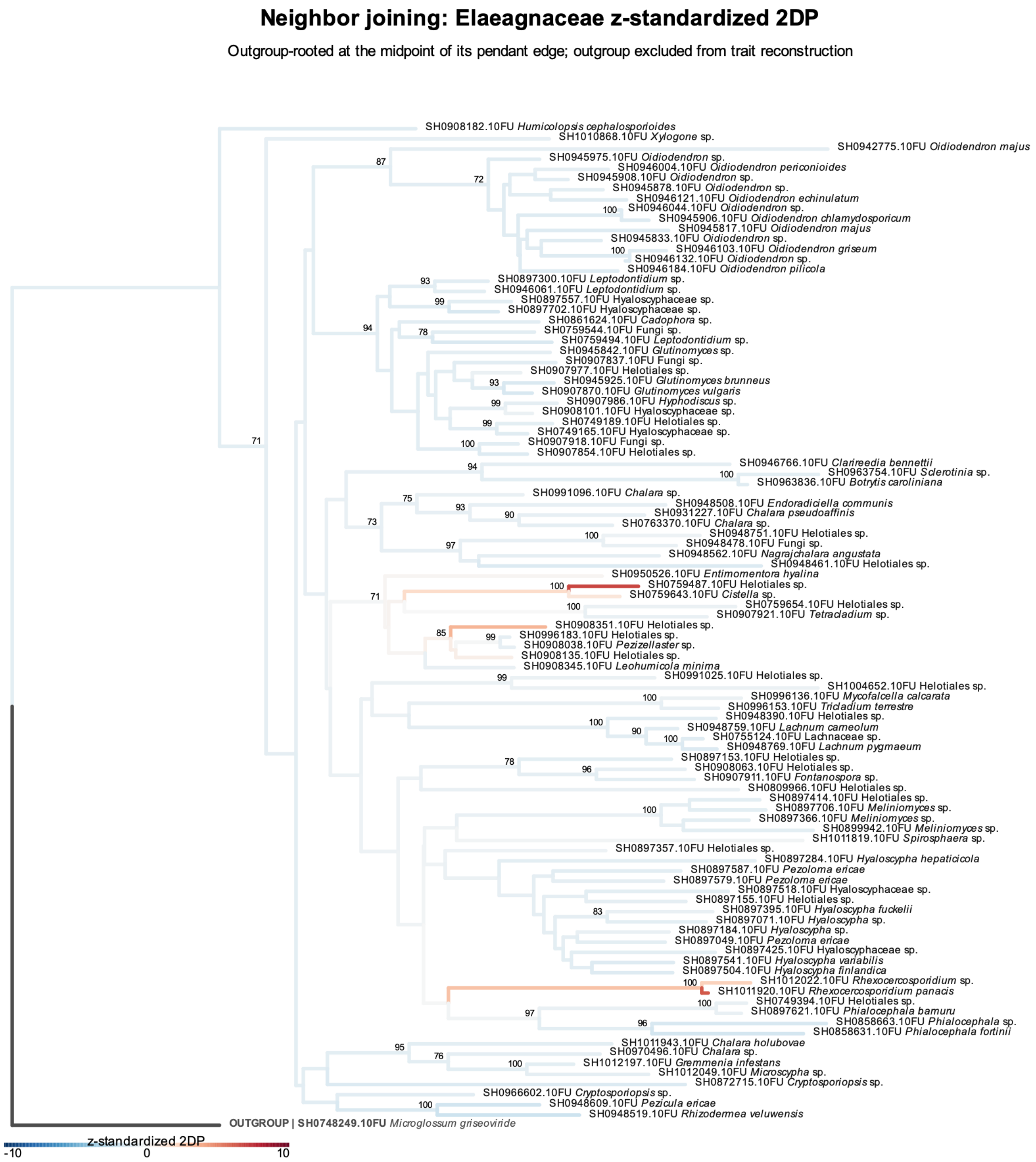


Figure S20: Phylogenetic patterns in the specificity of Helotiales fungi for Elaeagnaceae. The $z$-standardized 2DP values representing the specificity of individual Helotiales SHs for Elaeagnaceae are reflected by branch colors in the neighbor-joining phylogeny. Branch colors are scaled between $-10$ and 10 to emphasize variation around the significance thresholds. Bootstrap values greater than 70% are shown. See Table S1 for the results of Pagel's $\lambda$ analysis of phylogenetic conservatism.

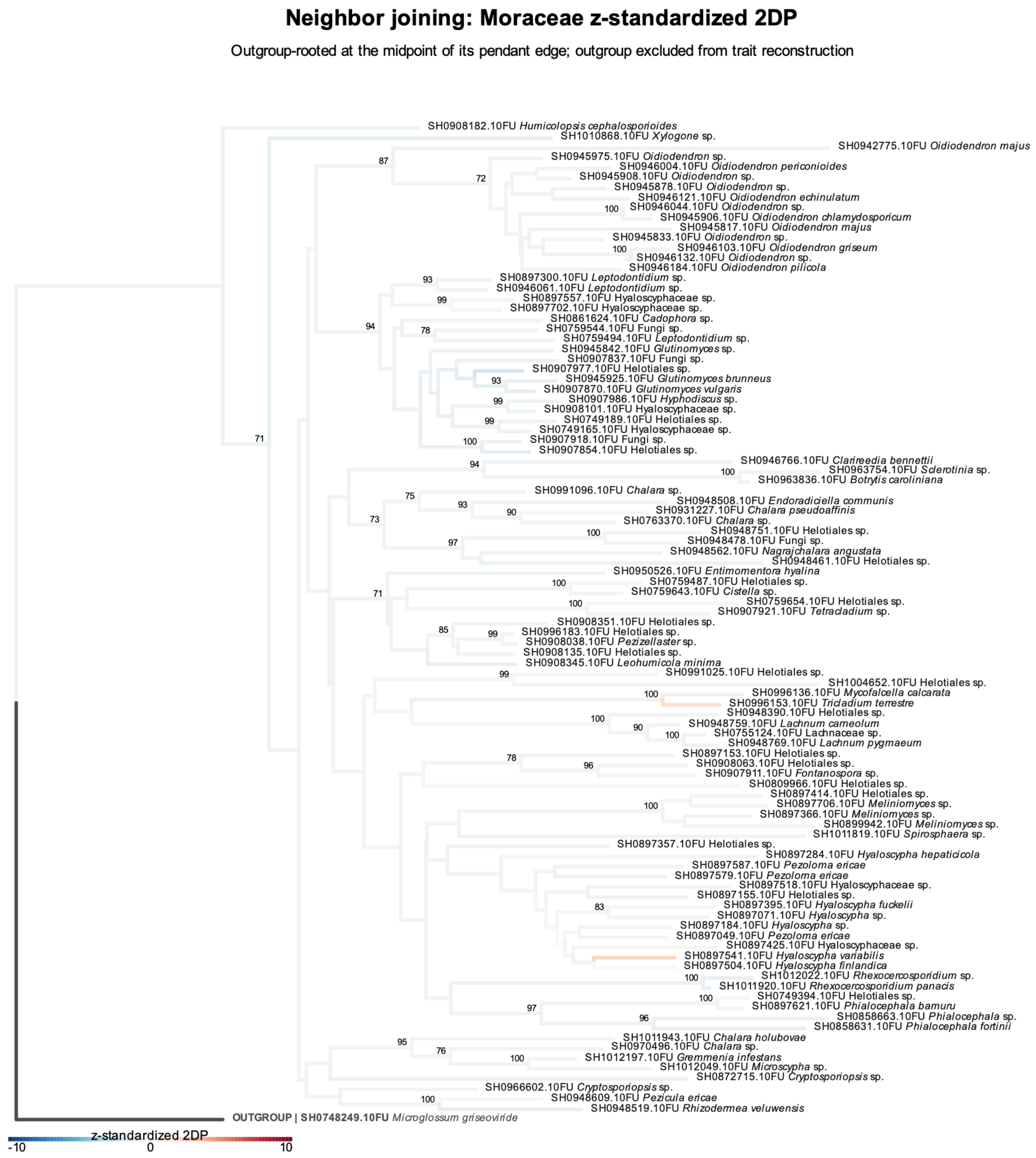


Figure S21: Phylogenetic patterns in the specificity of Helotiales fungi for Moraceae. The $z$-standardized 2DP values representing the specificity of individual Helotiales SHs for Moraceae are reflected by branch colors in the neighbor-joining phylogeny. Branch colors are scaled between −10 and 10 to emphasize variation around the significance thresholds. Bootstrap values greater than 70% are shown. See Table S1 for the results of Pagel's $\lambda$ analysis of phylogenetic conservatism.

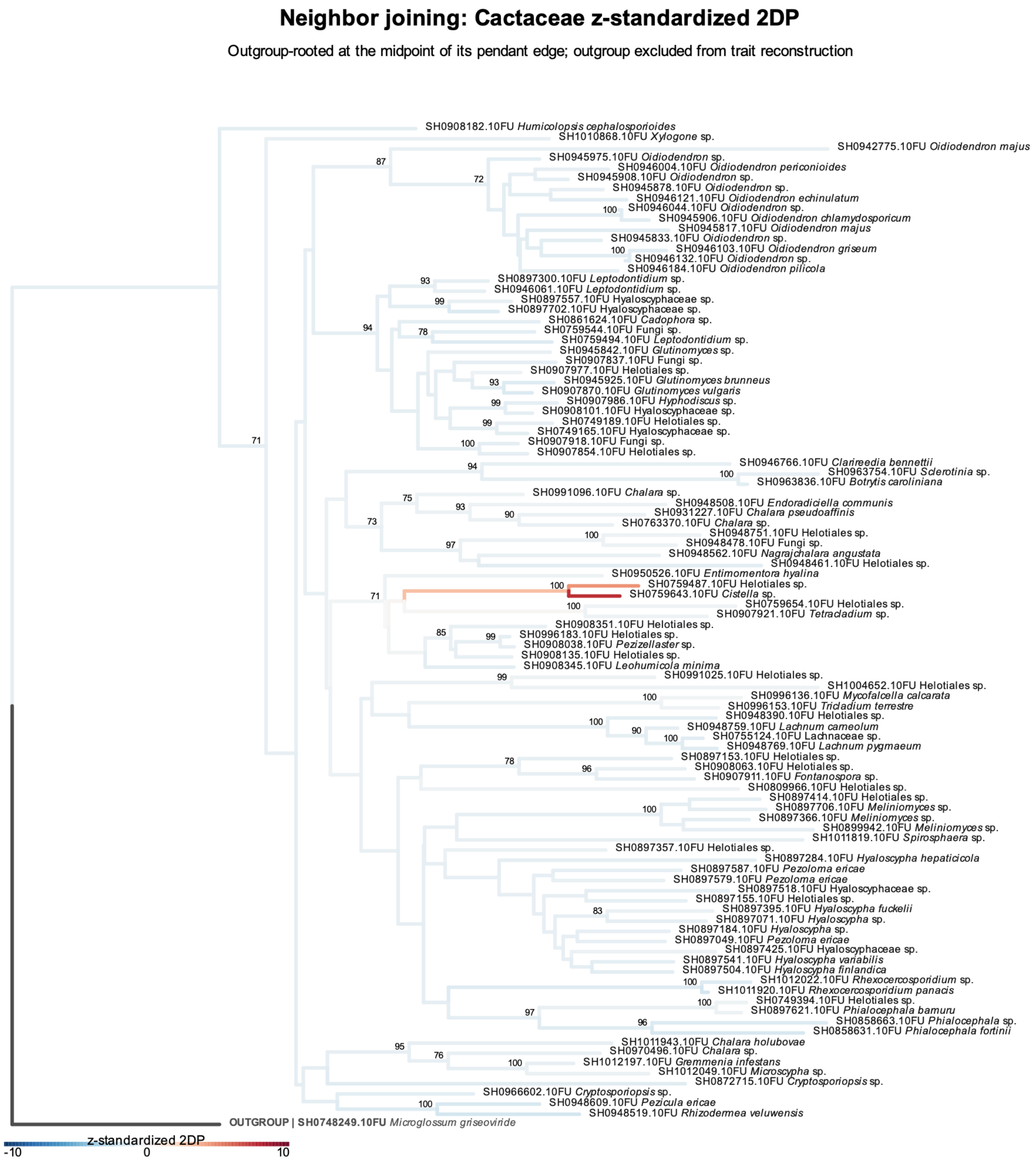


Figure S22: Phylogenetic patterns in the specificity of Helotiales fungi for Cactaceae. The $z$-standardized 2DP values representing the specificity of individual Helotiales SHs for Cactaceae are reflected by branch colors in the neighbor-joining phylogeny. Branch colors are scaled between −10 and 10 to emphasize variation around the significance thresholds. Bootstrap values greater than 70% are shown. See Table S1 for the results of Pagel's $\lambda$ analysis of phylogenetic conservatism.

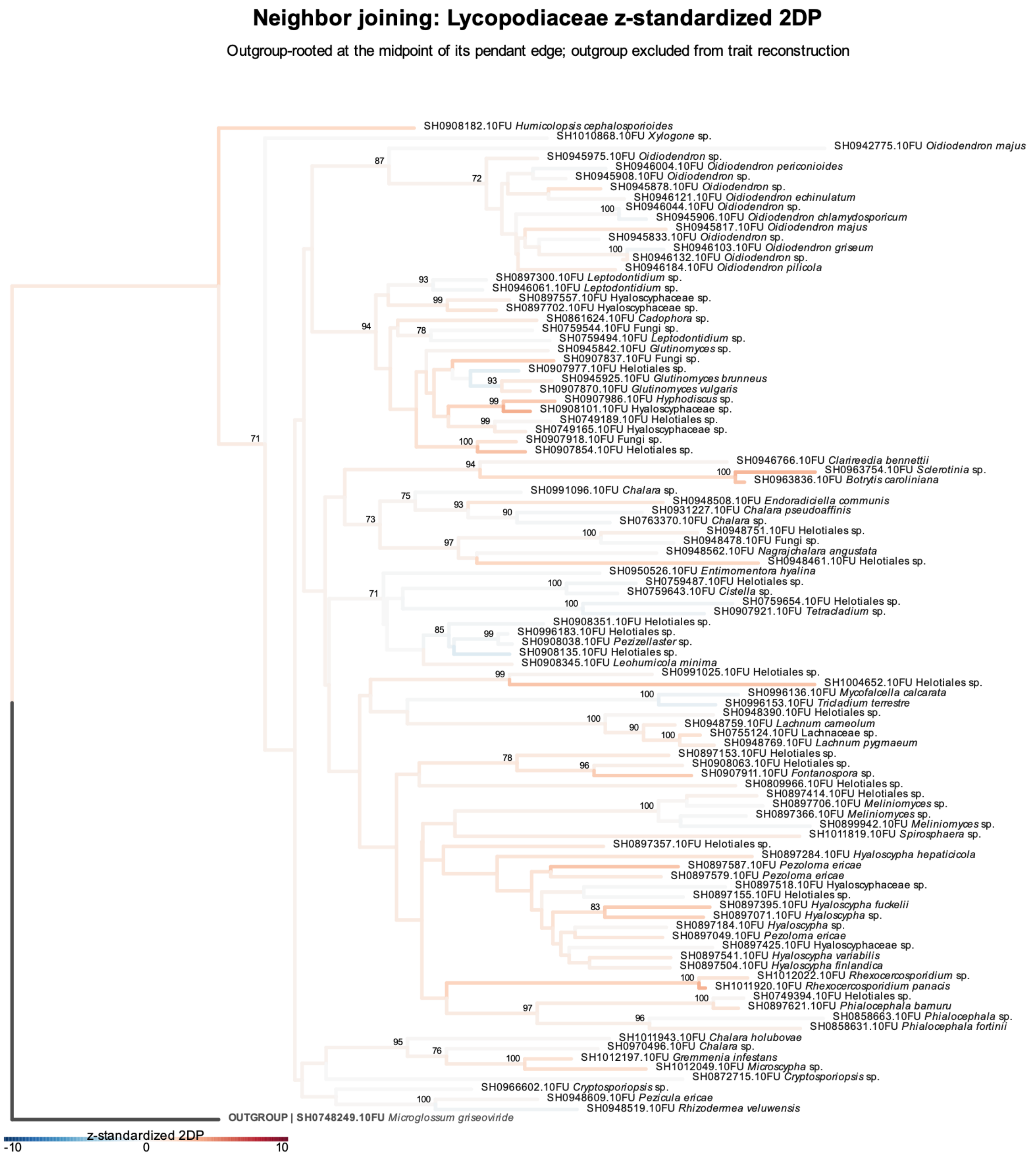


Figure S23: Phylogenetic patterns in the specificity of Helotiales fungi for Lycopodiaceae. The *z*-standardized 2DP values representing the specificity of individual Helotiales SHs for Lycopodiaceae are reflected by branch colors in the neighbor-joining phylogeny. Branch colors are scaled between −10 and 10 to emphasize variation around the significance thresholds. Bootstrap values greater than 70% are shown. See Table S1 for the results of Pagel's $\lambda$ analysis of phylogenetic conservatism.

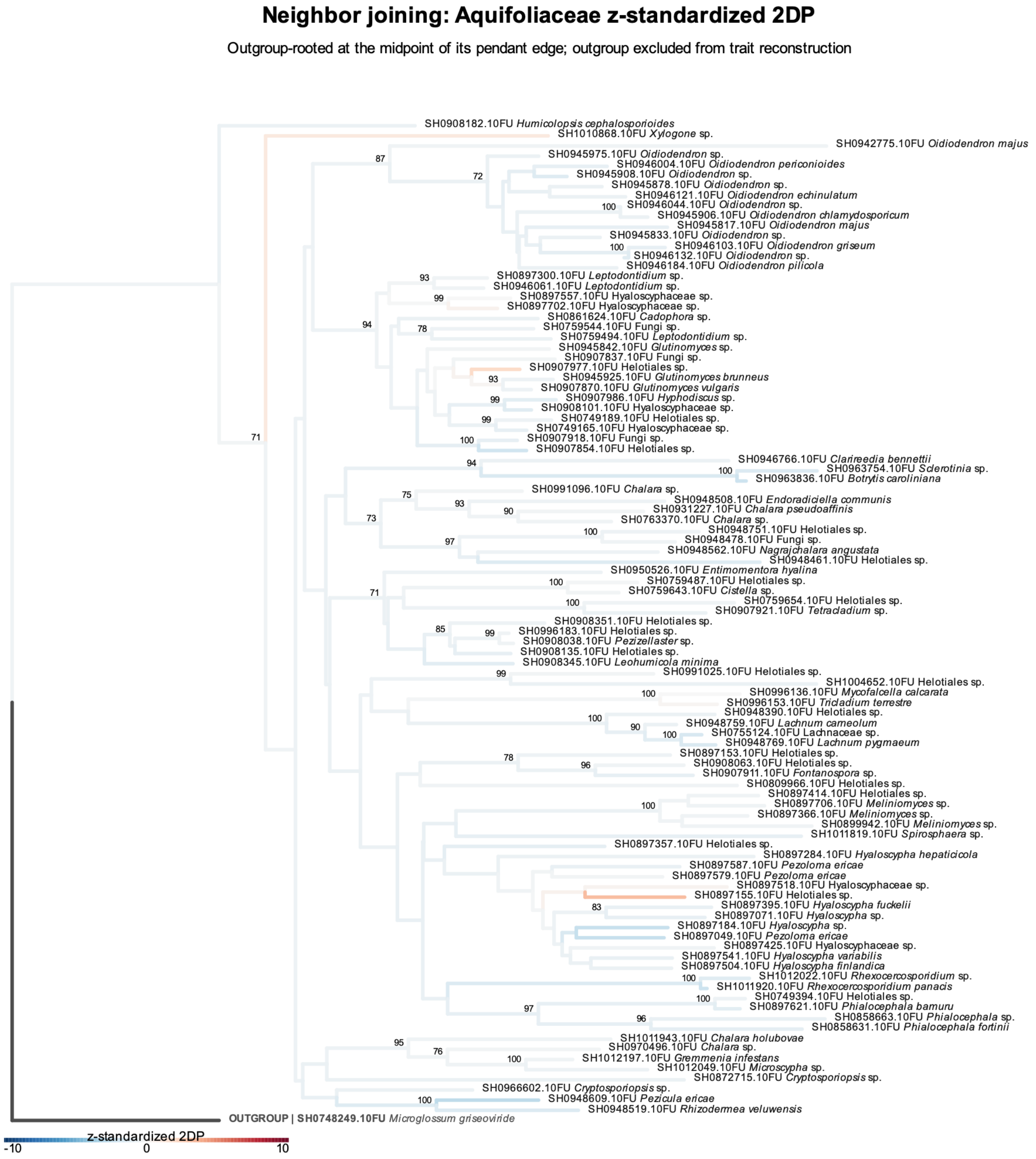


Figure S24: Phylogenetic patterns in the specificity of Helotiales fungi for Aquifoliaceae. The $z$-standardized 2DP values representing the specificity of individual Helotiales SHs for Aquifoliaceae are reflected by branch colors in the neighbor-joining phylogeny. Branch colors are scaled between −10 and 10 to emphasize variation around the significance thresholds. Bootstrap values greater than 70% are shown. See Table S1 for the results of Pagel's $\lambda$ analysis of phylogenetic conservatism.

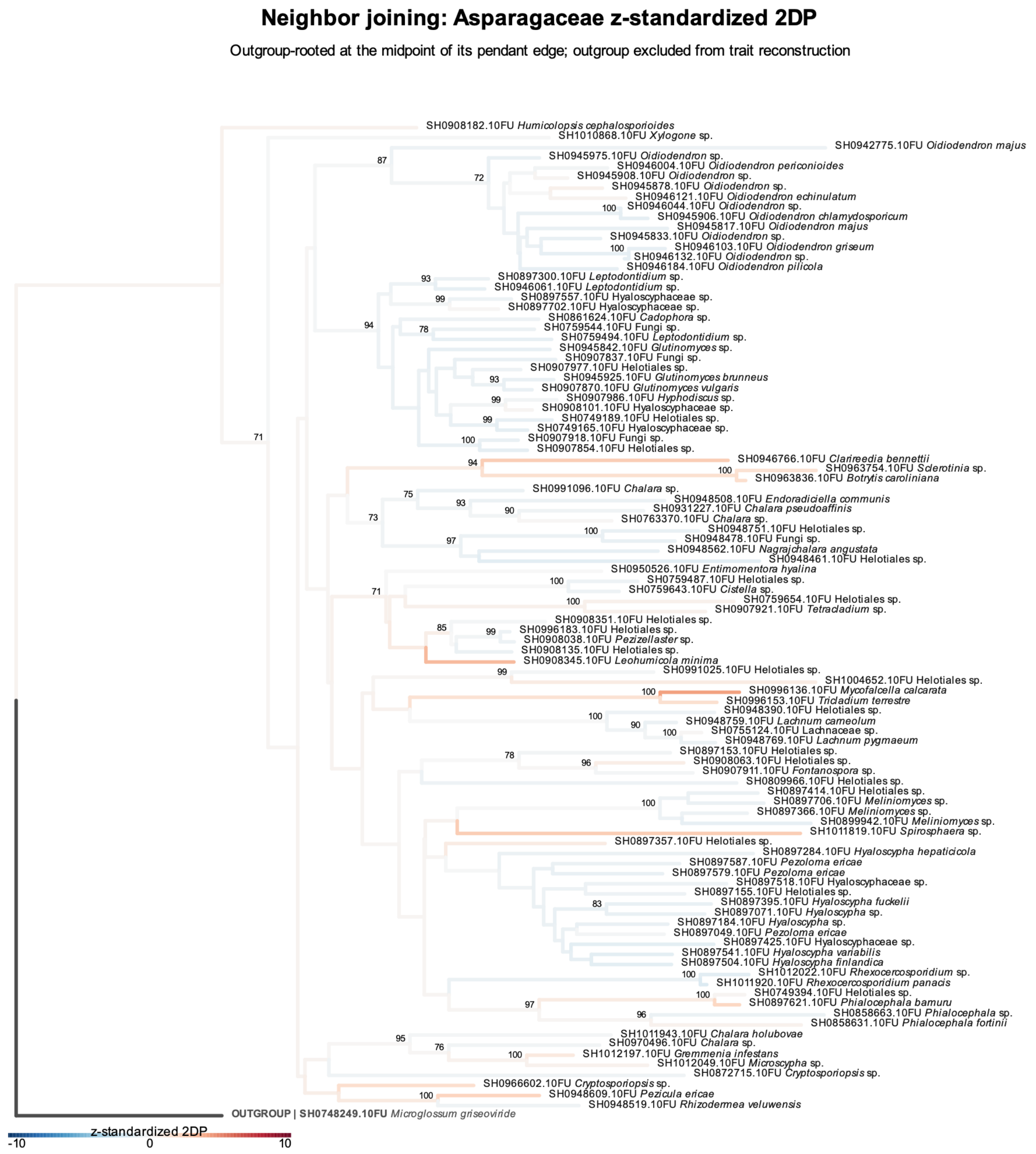


Figure S25: Phylogenetic patterns in the specificity of Helotiales fungi for Asparagaceae. The $z$-standardized 2DP values representing the specificity of individual Helotiales SHs for Asparagaceae are reflected by branch colors in the neighbor-joining phylogeny. Branch colors are scaled between −10 and 10 to emphasize variation around the significance thresholds. Bootstrap values greater than 70% are shown. See Table S1 for the results of Pagel's $\lambda$ analysis of phylogenetic conservatism.

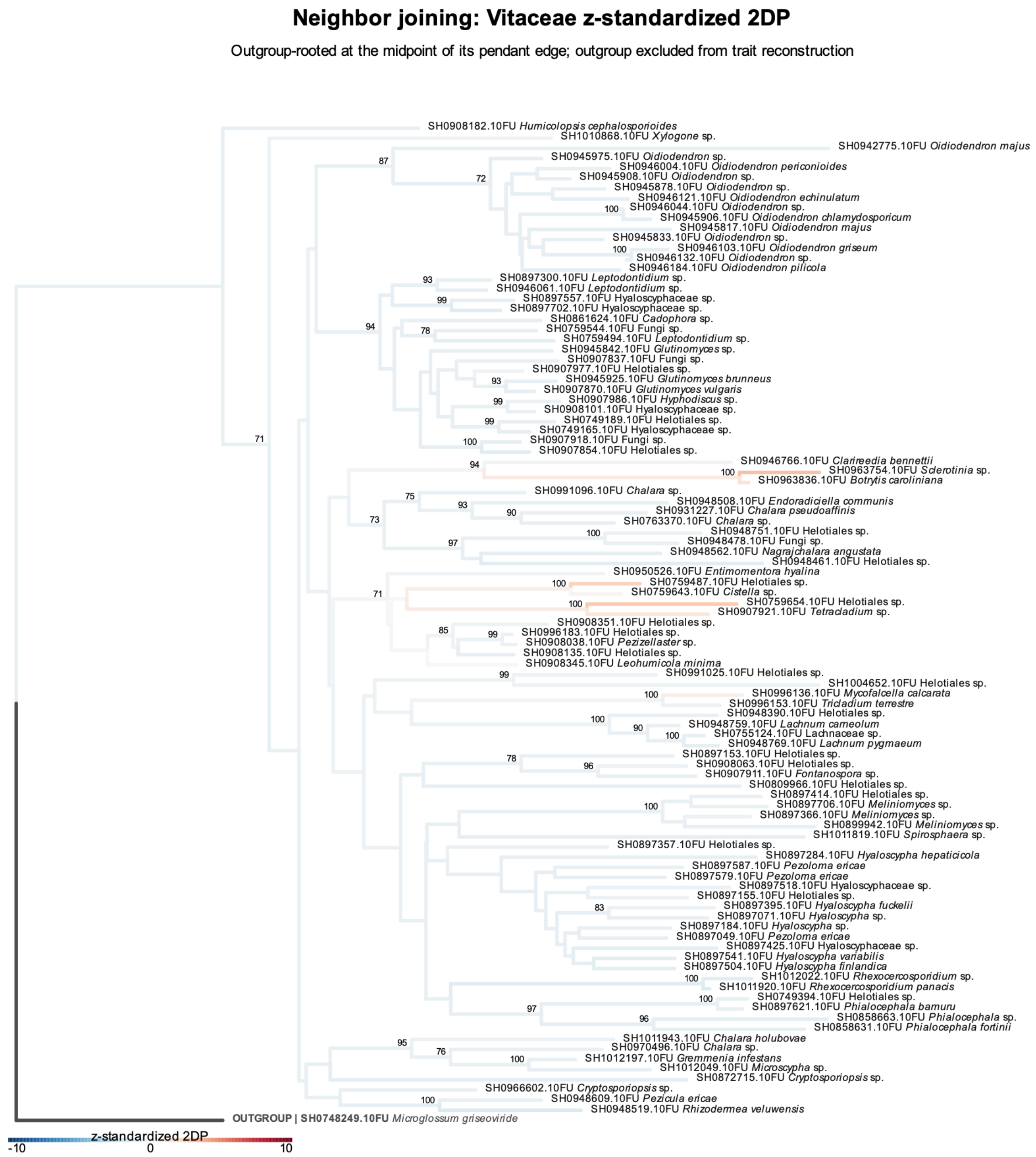


Figure S26: Phylogenetic patterns in the specificity of Helotiales fungi for Vitaceae. The *z*-standardized 2DP values representing the specificity of individual Helotiales SHs for Vitaceae are reflected by branch colors in the neighbor-joining phylogeny. Branch colors are scaled between −10 and 10 to emphasize variation around the significance thresholds. Bootstrap values greater than 70% are shown. See Table S1 for the results of Pagel's $\lambda$ analysis of phylogenetic conservatism.

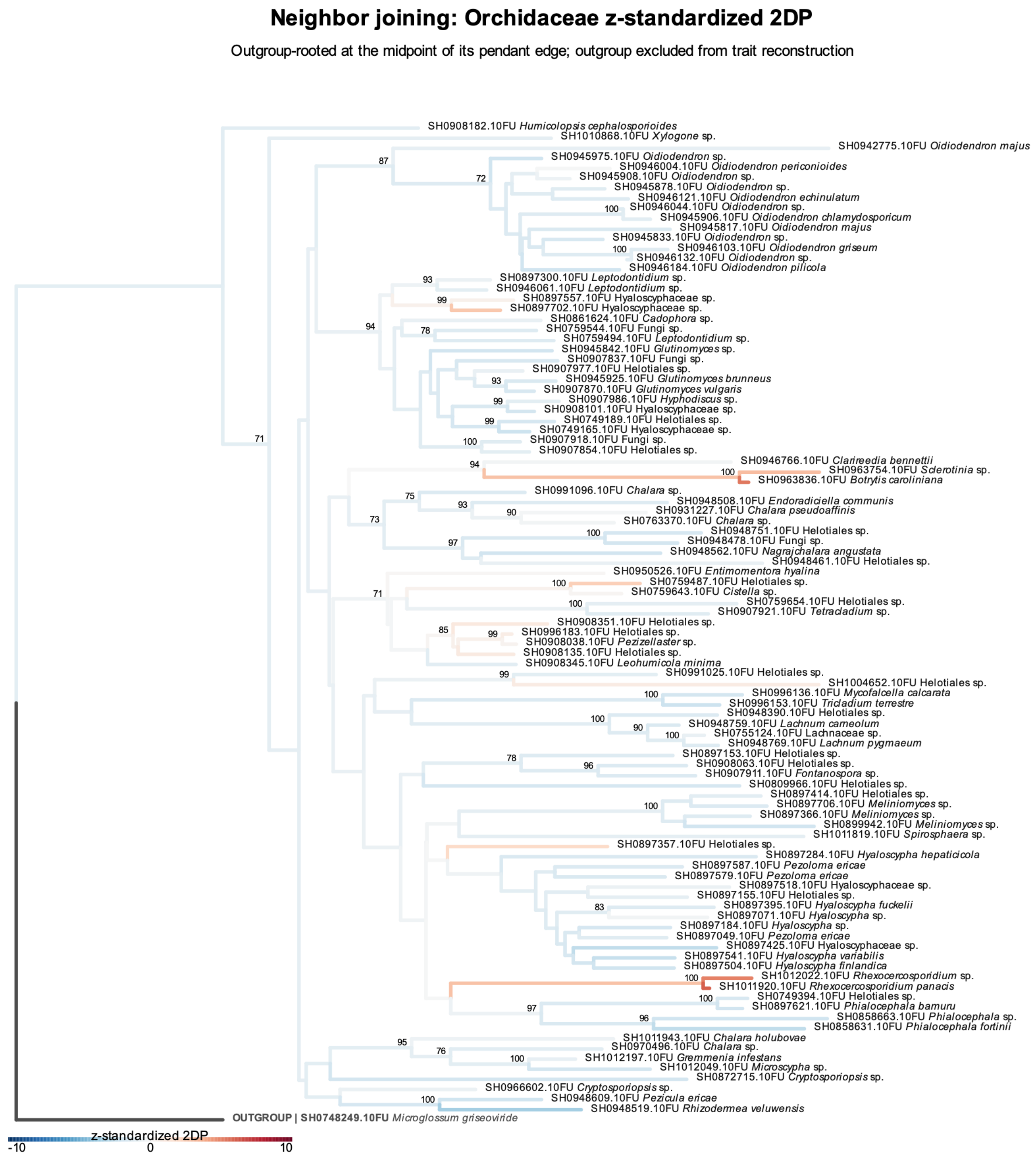

Figure S27: Phylogenetic patterns in the specificity of Helotiales fungi for Orchidaceae. The *z*-standardized 2DP values representing the specificity of individual Helotiales SHs for Orchidaceae are reflected by branch colors in the neighbor-joining phylogeny. Branch colors are scaled between −10 and 10 to emphasize variation around the significance thresholds. Bootstrap values greater than 70% are shown. See Table S1 for the results of Pagel's $\lambda$ analysis of phylogenetic conservatism.

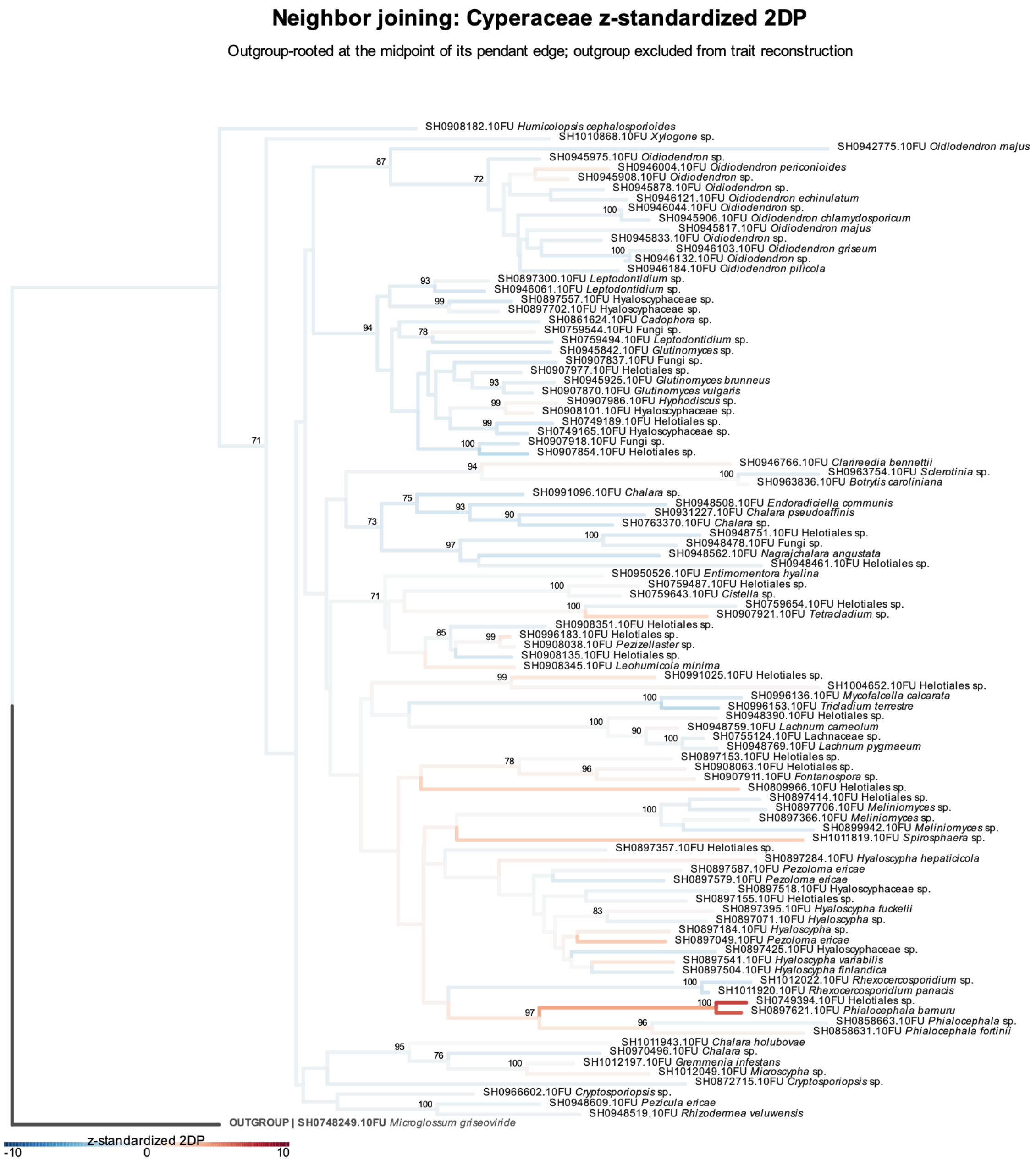


Figure S28: Phylogenetic patterns in the specificity of Helotiales fungi for Cyperaceae. The $z$-standardized 2DP values representing the specificity of individual Helotiales SHs for Cyperaceae are reflected by branch colors in the neighbor-joining phylogeny. Branch colors are scaled between −10 and 10 to emphasize variation around the significance thresholds. Bootstrap values greater than 70% are shown. See Table S1 for the results of Pagel's $\lambda$ analysis of phylogenetic conservatism.

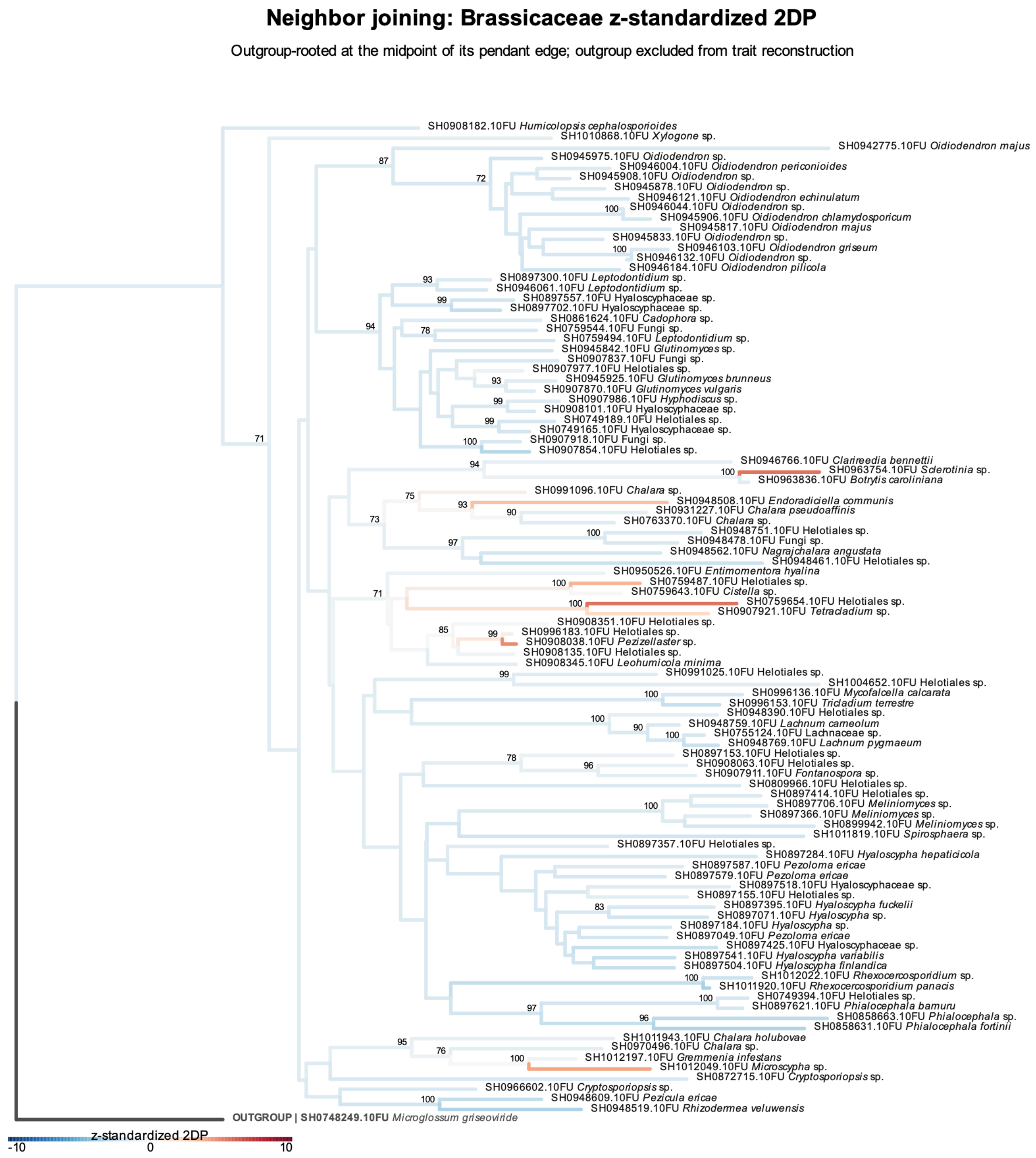


Figure S29: Phylogenetic patterns in the specificity of Helotiales fungi for Brassicaceae. The *z*-standardized 2DP values representing the specificity of individual Helotiales SHs for Brassicaceae are reflected by branch colors in the neighbor-joining phylogeny. Branch colors are scaled between −10 and 10 to emphasize variation around the significance thresholds. Bootstrap values greater than 70% are shown. See Table S1 for the results of Pagel's $\lambda$ analysis of phylogenetic conservatism.

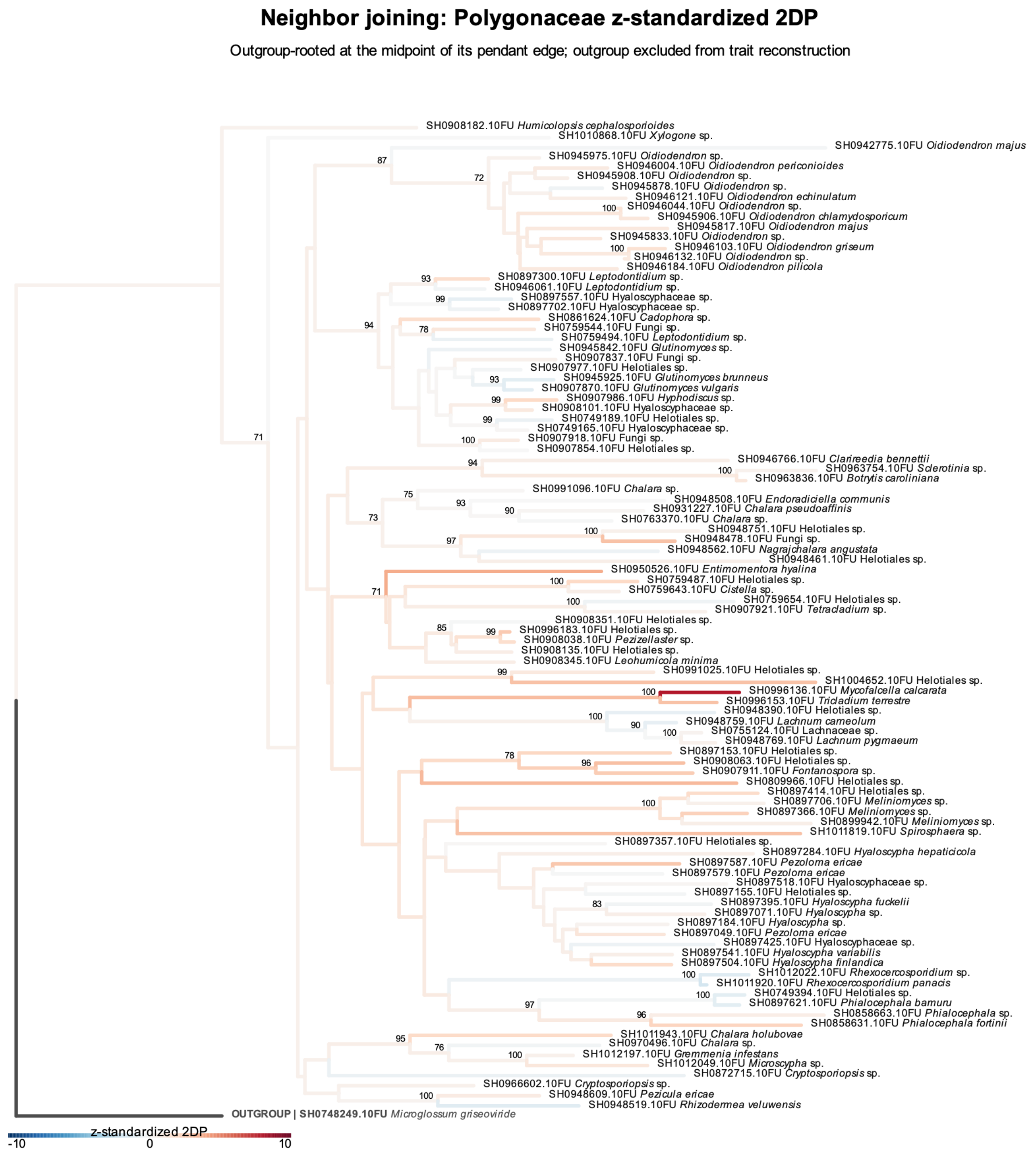


Figure S30: Phylogenetic patterns in the specificity of Helotiales fungi for Polygonaceae. The $z$-standardized 2DP values representing the specificity of individual Helotiales SHs for Polygonaceae are reflected by branch colors in the neighbor-joining phylogeny. Branch colors are scaled between −10 and 10 to emphasize variation around the significance thresholds. Bootstrap values greater than 70% are shown. See Table S1 for the results of Pagel's $\lambda$ analysis of phylogenetic conservatism.

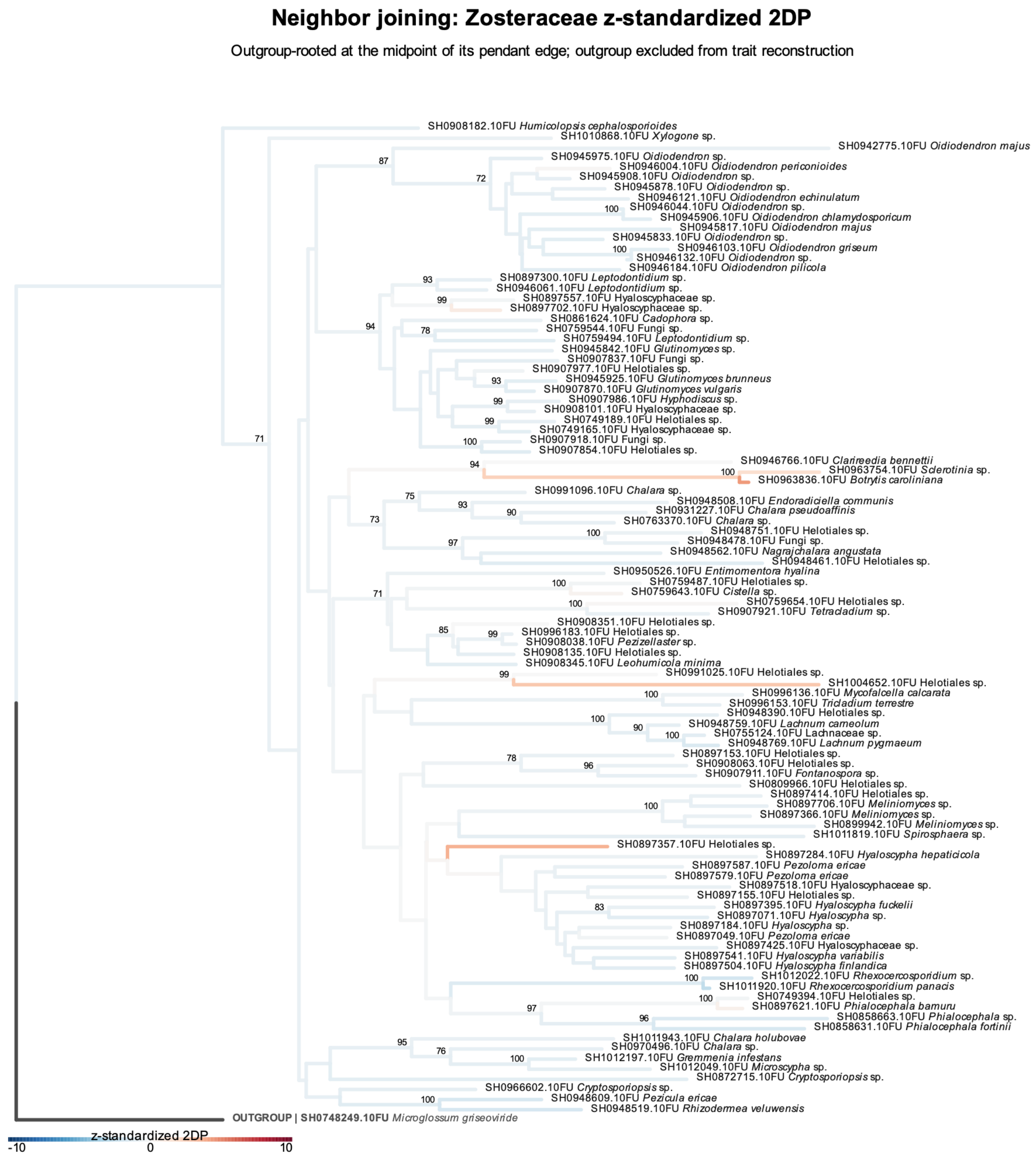


Figure S31: Phylogenetic patterns in the specificity of Helotiales fungi for Zosteraceae. The $z$-standardized 2DP values representing the specificity of individual Helotiales SHs for Zosteraceae are reflected by branch colors in the neighbor-joining phylogeny. Branch colors are scaled between −10 and 10 to emphasize variation around the significance thresholds. Bootstrap values greater than 70% are shown. See Table S1 for the results of Pagel's $\lambda$ analysis of phylogenetic conservatism.

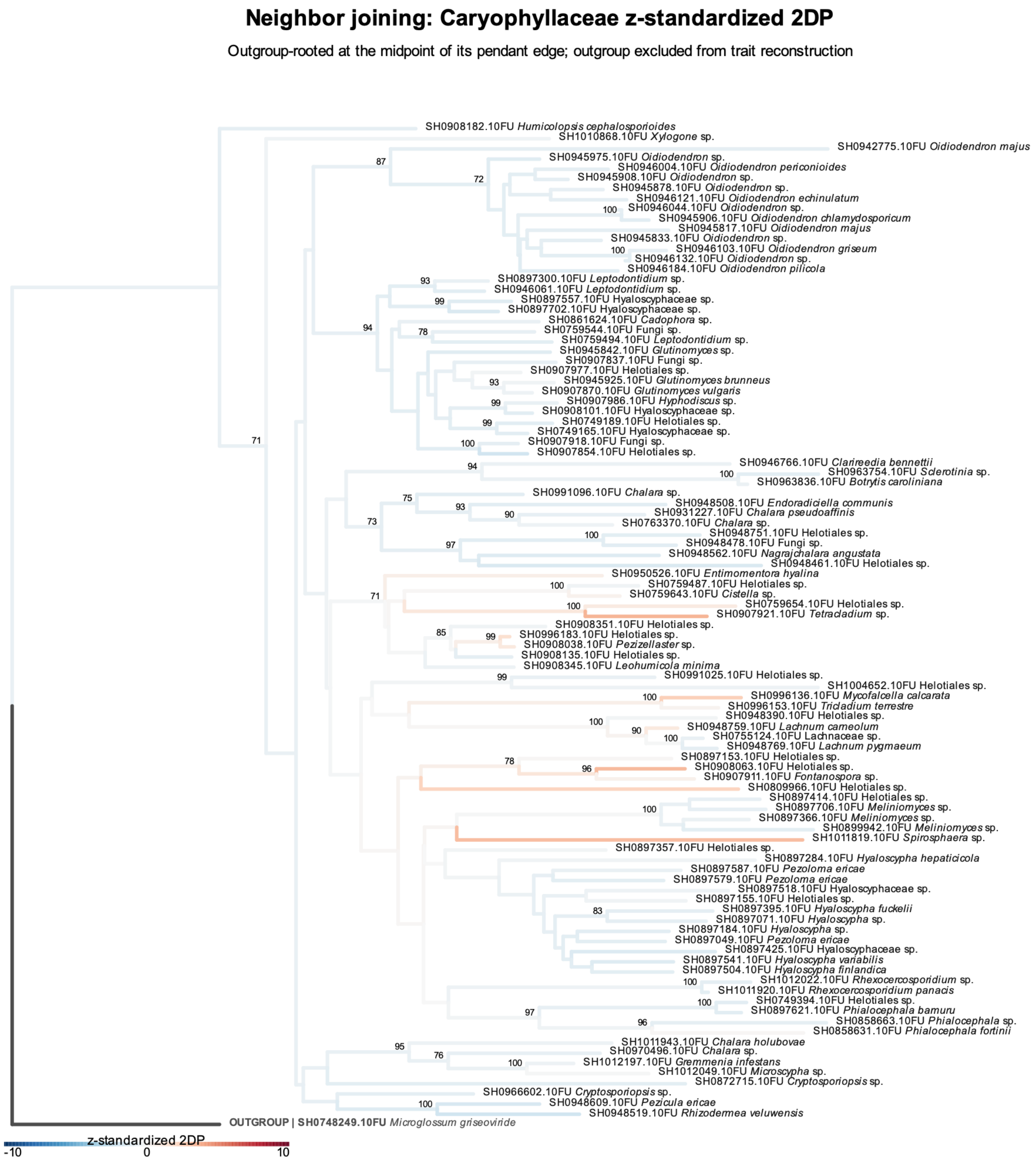


Figure S32: Phylogenetic patterns in the specificity of Helotiales fungi for Caryophyllaceae. The $z$-standardized 2DP values representing the specificity of individual Helotiales SHs for Caryophyllaceae are reflected by branch colors in the neighbor-joining phylogeny. Branch colors are scaled between −10 and 10 to emphasize variation around the significance thresholds. Bootstrap values greater than 70% are shown. See Table S1 for the results of Pagel's $\lambda$ analysis of phylogenetic conservatism.

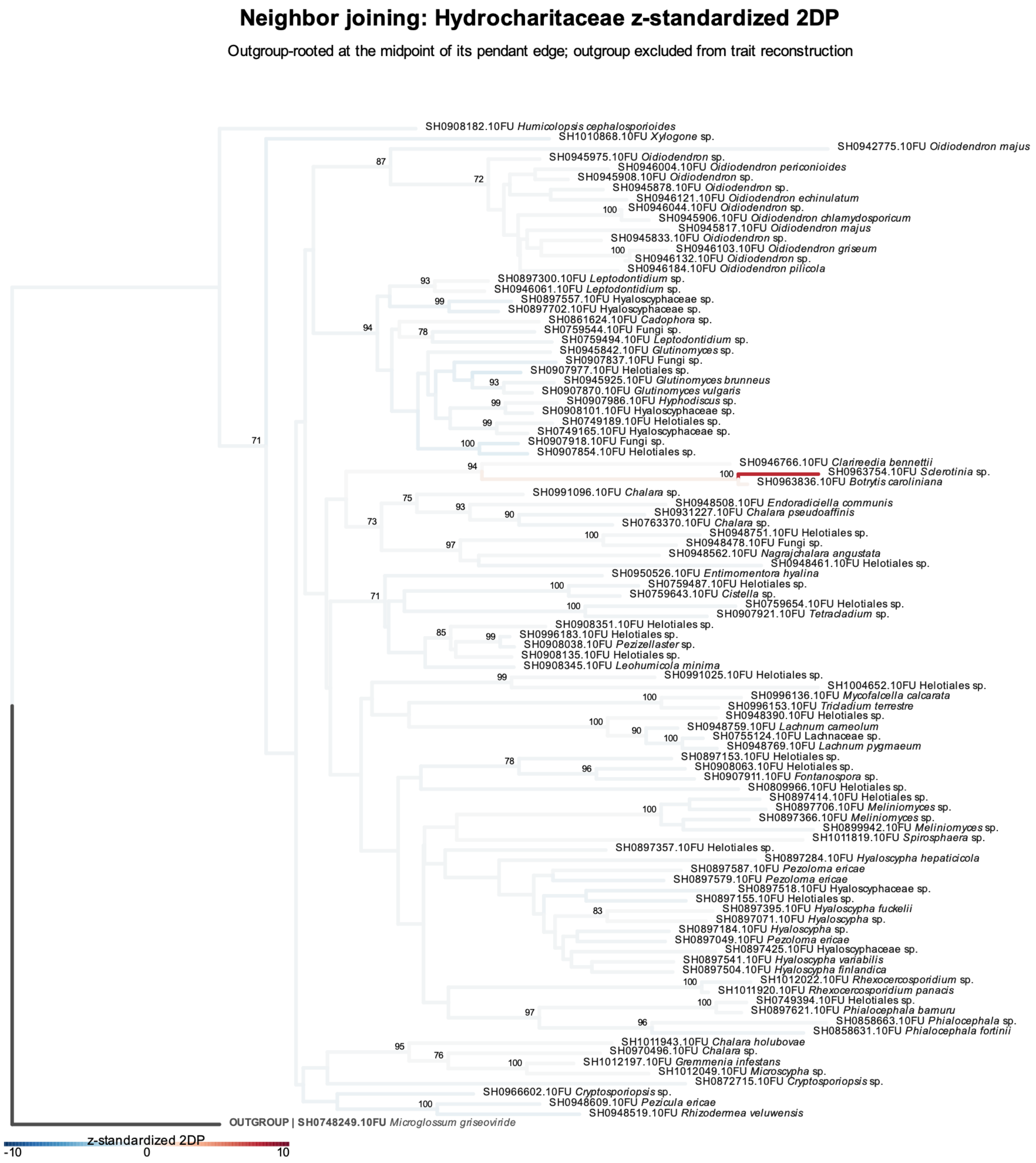


Figure S33: Phylogenetic patterns in the specificity of Helotiales fungi for Hydrocharitaceae. The *z*-standardized 2DP values representing the specificity of individual Helotiales SHs for Hydrocharitaceae are reflected by branch colors in the neighbor-joining phylogeny. Branch colors are scaled between −10 and 10 to emphasize variation around the significance thresholds. Bootstrap values greater than 70% are shown. See Table S1 for the results of Pagel's $\lambda$ analysis of phylogenetic conservatism.

Table S1: Phylogenetic conservatism in the association specificity for each plant family. The Pagel's λ statistics representing the phylogenetic conservatism of Helotiales fungi in their associations with each plant family (calculated based on *z*-standardized 2DP values) is shown with false discovery rate (FDR). Benjamini-Hochberg adjustment of *P* values was applied to the results of each phylogenetic method (maximum likelihood or neighbor joining).

| Phylogenetic method | Plant family | Pagel's λ | FDR |
|---|---|---|---|
| Neighbor joining | Fagaceae | 0.248 | 0.4593 |
| | Poaceae | 1.000 | < 0.0001 |
| | Ericaceae | 0.126 | 0.6293 |
| | Bromeliaceae | 0.000 | 1.0000 |
| | Betulaceae | 0.559 | 0.3595 |
| | Pinaceae | 0.701 | < 0.0001 |
| | Juglandaceae | 0.000 | 1.0000 |
| | Plantaginaceae | 0.398 | 0.0693 |
| | Orchidaceae | 0.751 | 0.0084 |
| | Rubiaceae | 0.169 | 0.9153 |
| | Cyperaceae | 0.419 | 0.1856 |
| | Brassicaceae | 0.268 | 0.1310 |
| | Asteraceae | 0.444 | 0.1310 |
| | Burmanniaceae | 0.000 | 1.0000 |
| | Rosaceae | 0.491 | 0.0147 |
| | Fabaceae | 0.812 | < 0.0001 |
| | Gentianaceae | 0.000 | 1.0000 |
| | Salicaceae | 0.640 | 0.0119 |
| | Polygonaceae | 0.000 | 1.0000 |
| | Myrtaceae | 0.000 | 1.0000 |
| | Zosteraceae | 0.000 | 1.0000 |
| | Elaeagnaceae | 0.167 | 0.6293 |
| | Caryophyllaceae | 0.565 | 0.0359 |
| | Moraceae | 0.000 | 1.0000 |
| | Cactaceae | 0.860 | < 0.0001 |
| | Lycopodiaceae | 0.103 | 1.0000 |
| | Aquifoliaceae | 0.000 | 1.0000 |
| | Asparagaceae | 0.430 | 0.5033 |
| | Vitaceae | 0.604 | 0.0008 |
| | Hydrocharitaceae | 0.000 | 1.0000 |

| Phylogenetic method | Plant family | Pagel's λ | FDR |
|---|---|---|---|
| Maximum likelihood | Fagaceae | 0.257 | 0.2294 |
| | Poaceae | 0.909 | < 0.0001 |
| | Ericaceae | 0.117 | 0.5754 |
| | Bromeliaceae | 0.000 | 1.0000 |
| | Betulaceae | 0.375 | 0.9130 |
| | Pinaceae | 0.678 | 0.0002 |
| | Juglandaceae | 0.000 | 1.0000 |
| | Plantaginaceae | 0.375 | 0.0960 |
| | Orchidaceae | 0.782 | 0.0057 |
| | Rubiaceae | 0.000 | 1.0000 |
| | Cyperaceae | 0.205 | 0.2275 |
| | Brassicaceae | 0.336 | 0.1582 |
| | Asteraceae | 0.422 | 0.1628 |
| | Burmanniaceae | 0.000 | 1.0000 |
| | Rosaceae | 0.462 | 0.0563 |
| | Fabaceae | 0.806 | 0.0001 |
| | Gentianaceae | 0.000 | 1.0000 |
| | Salicaceae | 0.609 | 0.0304 |
| | Polygonaceae | 0.000 | 1.0000 |
| | Myrtaceae | 0.000 | 1.0000 |
| | Zosteraceae | 0.000 | 1.0000 |
| | Elaeagnaceae | 0.215 | 0.4422 |
| | Caryophyllaceae | 0.559 | 0.0063 |
| | Moraceae | 0.000 | 1.0000 |
| | Cactaceae | 0.812 | 0.0004 |
| | Lycopodiaceae | 0.368 | 0.2673 |
| | Aquifoliaceae | 0.000 | 1.0000 |
| | Asparagaceae | 0.437 | 0.4422 |
| | Vitaceae | 0.626 | 0.0005 |
| | Hydrocharitaceae | 0.000 | 1.0000 |
